# ArchR: An integrative and scalable software package for single-cell chromatin accessibility analysis

**DOI:** 10.1101/2020.04.28.066498

**Authors:** Jeffrey M. Granja, M. Ryan Corces, Sarah E. Pierce, S. Tansu Bagdatli, Hani Choudhry, Howard Y. Chang, William J. Greenleaf

## Abstract

The advent of large-scale single-cell chromatin accessibility profiling has accelerated our ability to map gene regulatory landscapes, but has outpaced the development of robust, scalable software to rapidly extract biological meaning from these data. Here we present a software suite for single-cell <u>a</u>nalysis of regulatory <u>ch</u>romatin in <u>R</u> (ArchR; www.ArchRProject.com) that enables fast and comprehensive analysis of single-cell chromatin accessibility data. ArchR provides an intuitive, user-focused interface for complex single-cell analyses including doublet removal, single-cell clustering and cell type identification, robust peak set generation, cellular trajectory identification, DNA element to gene linkage, transcription factor footprinting, mRNA expression level prediction from chromatin accessibility, and multi-omic integration with scRNA-seq. Enabling the analysis of over 1.2 million single cells within 8 hours on a standard Unix laptop, ArchR is a comprehensive analytical suite for end-to-end analysis of single-cell chromatin accessibility data that will accelerate the understanding of gene regulation at the resolution of individual cells.

## INTRODUCTION

Single-cell approaches have revolutionized our understanding of biology, opening the door for a wide array of applications ranging from interrogation of cellular heterogeneity to identification of disease-specific processes. The advent of single-cell approaches for the assay for transposase-accessible chromatin using sequencing (scATAC-seq) has made it possible to study chromatin accessibility and gene regulation in single cells^1, 2^. These chromatin-based assays have illuminated cell type-specific biology and provided insights into complex biological processes previously hidden by ensemble averaging^3–7^. Recent methodological advances have increased the throughput of scATAC-seq, enabling a single lab to generate data from hundreds of thousands of cells on the timescale of weeks^5, 6, 8^. These advances have been driven by an increased interest in chromatin-based gene regulation across a diversity of cellular contexts and biological systems^1, 2, 5, 6, 8^. This capacity for data generation has outpaced the development of intuitive, robust, and comprehensive software for analysis of these scATAC-seq datasets^9^ – a crucial requirement that would facilitate the broad utilization of these methods of investigating gene regulation at cellular resolution.

To this end, we sought to develop a user-oriented software suite for both routine and advanced analysis of massive-scale single-cell chromatin accessibility data from diverse sources without the need for high-performance computing environments. This package for single-cell Analysis of Regulatory Chromatin in R (ArchR; www.ArchRProject.com) provides a facile platform to interrogate scATAC-seq data from multiple scATAC-seq implementations, including the 10x Genomics Chromium system^6, 7^, the Bio-Rad droplet scATAC-seq system^8^, single-cell combinatorial indexing^2, 5^, and the Fluidigm C1 system^1, 4^ (**Fig. 1a**). ArchR provides a user-focused interface for complex scATAC-seq analysis such as marker feature identification, transcription factor (TF) footprinting, an interactive genome browsing, scRNA-seq integration, and cellular trajectory analysis (**Fig. 1a**). When compared to other existing tools, such as SnapATAC^10^ and Signac^11^, ArchR provides a more extensive set of features with substantially improved performance benchmarks (**Supplementary Fig. 1a**). Moreover, ArchR is designed to provide the speed and flexibility to support interactive analysis, enabling iterative extraction of meaningful biological interpretations.

**Figure 1.**
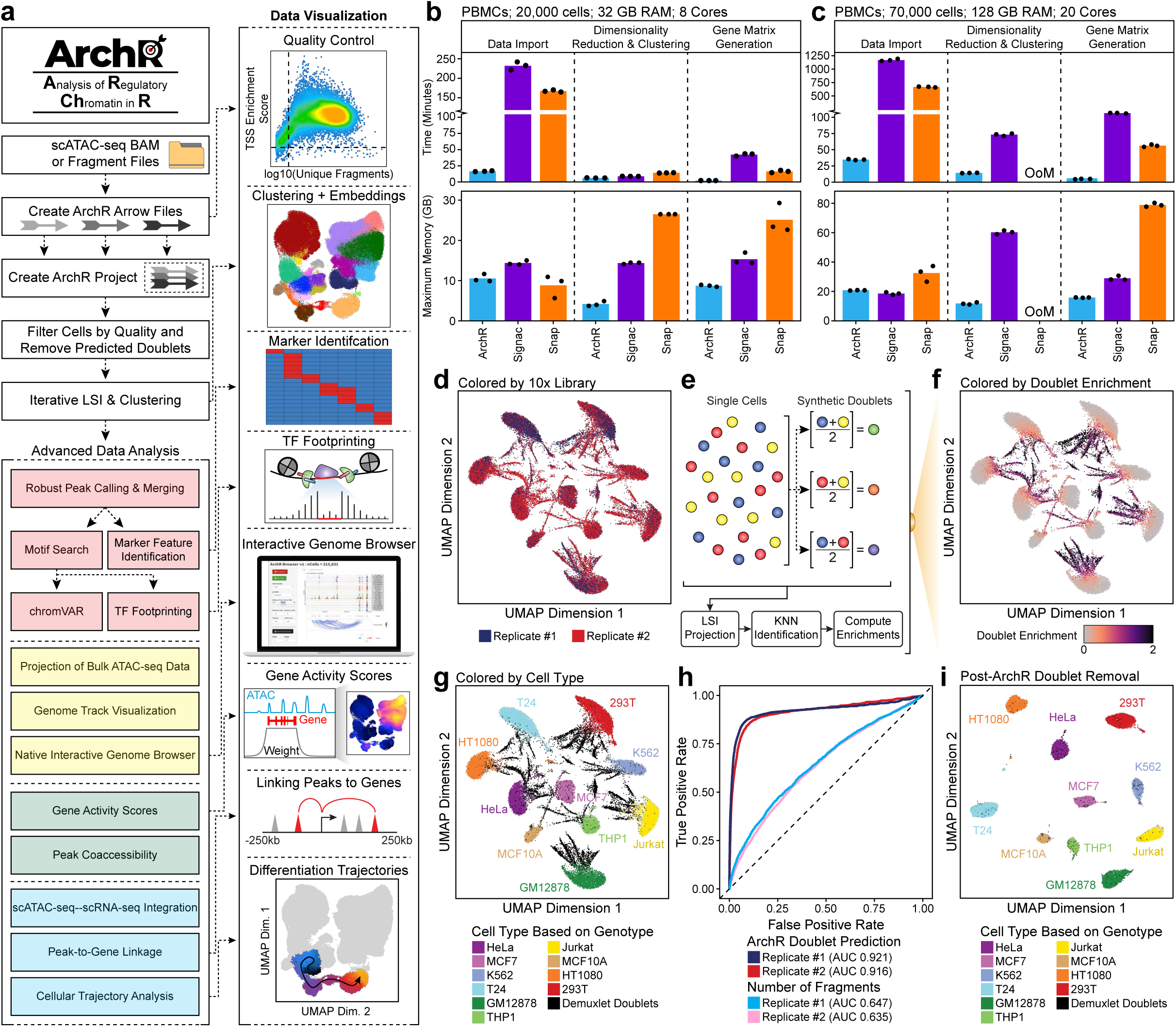
ArchR: A rapid, extensible, and comprehensive scATAC-seq analysis platform. **a.** Schematic of the ArchR workflow from input of pre-aligned scATAC-seq data as BAM or fragment files to diverse data analysis. **b-c.** Comparison of run time and memory usage by ArchR, Signac, and SnapATAC for the analysis of (**b**) ∼20,000 PBMC cells using 32 GB RAM and 8 cores or (**c**) ∼70,000 PBMC cells using 128 GB RAM and 20 cores. Dots represent individual replicates of benchmarking analysis. **d.** Initial UMAP embedding of scATAC-seq data from 2 replicates of the cell line mixing experiment (N = 38,072 total cells from 10 different cell lines) colored by replicate number. **e.** Schematic of doublet identification with ArchR. **f-g.** Initial UMAP embedding of scATAC-seq data from 2 replicates of the cell line mixing experiment (N = 38,072 total cells from 10 different cell lines) colored by (**f**) the enrichment of projected synthetic doublets or (**g**) the demuxlet identification labels based on genotype identification using SNPs within accessible chromatin sites. **h.** Receiver operating characteristic (ROC) curves of doublet prediction using synthetic doublet projection enrichment or the number of nuclear fragments per cell compared to demuxlet as a ground truth. The area under the curve (AUC) for these ROC curves are annotated below. **i.** UMAP after ArchR doublet removal of scATAC-seq data from 2 replicates of the cell line mixing experiment (N = 27,220 doublet-filtered cells from 10 different cell lines) colored by demuxlet identification labels based on genotype identification using SNPs within accessible chromatin sites.

## RESULTS

### The ArchR framework

ArchR takes as input aligned BAM or fragment files, which are first parsed in small chunks per chromosome, read in parallel to conserve memory, then efficiently stored on disk using the compressed random-access hierarchical data format version 5 (HDF5) file format. These HDF5 files form the constituent pieces of an ArchR analysis which we call “Arrow” files. Arrow files are grouped into an “ArchR Project”, a compressed R data file that is stored in memory, which provides an organized, rapid, and low memory-use framework for manipulation of the larger arrow files stored on disk (**Supplementary Fig. 1b**). Arrow files are always accessed in chunks using parallel read and write operations that minimize memory while efficiently using the multi-processor capabilities of most standard computers (**Supplementary Fig. 1c-d**). Moreover, the base file size of Arrow files remains smaller than the input fragment files across various cellular inputs (**Supplementary Fig. 2a-b**). These efficiencies provide substantial improvements in speed and memory usage compared to scATAC-seq software packages such as SnapATAC and Signac.

### ArchR enables efficient and comprehensive single-cell chromatin accessibility analysis

To benchmark the performance of ArchR, we collected three diverse publicly available datasets (**Supplementary Table 1**): (i) peripheral blood mononuclear cells (PBMCs) that represent discrete primary cell types^6, 7^ (**Supplementary Fig. 2c-e**), (ii) bone marrow stem/progenitor cells and differentiated cells that represent a continuous cellular hierarchy^7^ (**Supplementary Fig. 2f-h**), and (iii) a large atlas of murine cell types from diverse organ systems^5^ (**Supplementary Fig. 2i-k**). Prior to downstream analysis, we performed rigorous quality control of each dataset to remove low quality cells. To assess per-cell data quality, ArchR computes TSS enrichment scores, which have become the standard for bulk ATAC-seq analysis (https://www.encodeproject.org/atac-seq/) and provide clearer separation of low- and high-quality cells compared to other metrics such as the fraction of reads in promoters^10^ (**Supplementary Fig. 2c,f**).

To quantify the ability of ArchR to analyze large-scale data, we compared the performance of ArchR to that of SnapATAC and Signac for three of the major scATAC-seq analytical steps across these three datasets using two different computational infrastructures (**Supplementary Fig 3a and Supplementary Table 2**). We observed that ArchR outperforms SnapATAC and Signac in speed and memory usage across all comparisons, enabling analysis of 70,000 cell datasets in under and hour with 32 GB of RAM and 8 cores (**Fig. 1b-c and Supplementary Fig. 3b-i**). Additionally, when analyzing a 70,000-cell dataset, SnapATAC exceeded the available memory in the high memory setting (128 GB RAM, 20 cores) (**Fig. 1c**) and both SnapATAC and Signac exceeded the available memory in the low memory setting (32 GB RAM, 8 cores) (**Supplementary Fig. 3c**), while ArchR completed these analyses faster and without exceeding the available memory. Lastly, ArchR can analyze scATAC-seq data directly from BAM files, enabling the analysis of data from diverse single-cell platforms including the sci-ATAC-seq murine atlas^5^ (**Supplementary Fig. 3j-k**).

### ArchR identifies putative doublets in scATAC-seq data

The presence of so called “doublets” – two cells that are captured within the same nano-reaction (i.e. a droplet) and thus indexed with the same cellular barcode – often complicate single-cell analysis. Doublets appear as a superposition of signals from both cells, leading to the false appearance of distinct clusters or false connections between distinct cell types. To mitigate this issue, we designed a doublet detection and removal algorithm as part of ArchR. Similar to methods employed for doublet detection in scRNA-seq^12, 13^, ArchR identifies heterotypic doublets by bioinformatically generating a collection of synthetic doublets, projecting these synthetic doublets into the low-dimensional data embedding, then identifying the nearest neighbors to these synthetic doublets as doublets themselves^12, 13^ (**Fig. 1d-f**). To validate this approach, we carried out scATAC-seq on a mixture of 10 highly distinct human cell lines (N = 38,072 cells), allowing for genotype-based identification of doublets via demuxlet^14^ as a ground-truth comparison for computational identification of doublets by ArchR (**Fig. 1g and Supplementary Fig. 4a**). Using an unbiased optimization for the projection of synthetic doublets, we identified robust parameters **(Supplementary Fig. 4b**) for doublet prediction (ROC = 0.918) which significantly outperformed doublet prediction based on the total number of accessible fragments (ROC = 0.641) (**Fig. 1h and Supplementary Fig 4c-h**). With these predicted doublets excluded, the remaining cells formed 10 large groups according to their cell line of origin (**Fig. 1i**). ArchR’s implementation of heterotypic doublet elimination reduces false cluster identification and thus improves the fidelity of downstream results.

### ArchR provides high-resolution and efficient dimensionality reduction of scATAC-seq data

ArchR additionally provides methodological improvements over other available software. One of the most fundamental aspects of ATAC-seq analysis is the identification of a feature set (i.e. a peak set) for downstream analysis. In the context of single-cell ATAC-seq, identification of peak regions prior to cluster identification requires peak calling from all cells as a single merged group. This effectively obscures cell type-specific chromatin accessibility which distorts downstream analyses. For Signac, a counts matrix is created using a pre-determined peak set, preventing the contribution of peaks that are specific to lowly represented cell types. Instead of using a pre-determined peak set, SnapATAC creates a genome-wide tiled matrix of 5-kb bins, allowing for unbiased genome-wide identification of cell type-specific chromatin accessibility. However, 5-kb bins are substantially larger than the average regulatory element (∼300-500 bp containing TF binding sites less than 50 bp)^15–17^, thus causing multiple regulatory elements to be grouped together, again obscuring cell type-specific biology. To avoid both of these pitfalls, ArchR operates on a genome-wide tiled matrix of 500-bp bins, allowing for the sensitivity to capture cell type-specific biology at individual regulatory elements across the entire genome. Despite this 10-fold higher resolution tile matrix, ArchR stores both per-tile accessibility information and all ATAC-seq fragments in an Arrow file that is smaller than either the original input fragments or the Snap file from SnapATAC containing the genome-wide tiled matrix at only 5-kb resolution (**Supplementary Fig. 2a-b).**

One major application of single-cell analysis is the identification of cellular subsets through dimensionality reduction and clustering. For dimensionality reduction, ArchR uses an optimized iterative latent semantic indexing (LSI) method^6, 7^ (**Supplementary Fig. 5a**), Signac uses an LSI method, and SnapATAC uses a method based on Jaccard indices. When directly comparing the results from these different dimensionality reduction methods, ArchR identified similar clusters to other methods while being less biased by low-quality cells and doublets (**Supplementary Fig. 5b**). However, when comparing clustering of the bone marrow cell dataset, we found that ArchR alone maintained the continuous cellular hierarchy expected in this biological system **(Supplementary Fig. 6a).**

To enable the efficient examination of extremely large datasets, ArchR implements a novel estimated LSI dimensionality reduction by first creating an iterative LSI reduction from a subset of the total cells, then linearly projecting the remainder of cells into this reduced dimension space using LSI projection^7^ (**Supplementary Fig. 7a**). We compared this approach to the landmark diffusion map (LDM) estimation method used by SnapATAC which uses a non-linear reduction based on a subset of cells and then projects the remainder of the cells into this subspace using LDM projection. When comparing “landmark” subsets of different cell numbers, the estimated LSI approach implemented by ArchR was more consistent and could recapitulate the clusters called and the overall structure of the data with as few as 50 cells across both the PBMC (N = 27,845 cells) and bone marrow cell (N = 26,748 cells) datasets (**Supplementary Fig. 7b and 8a-b**). We speculate that this observed robustness stems from the linearity of the LSI projection as compared to LDM projection, which occurs in a non-linear subspace. The estimated LSI approach implemented by ArchR is also faster than the estimated LDM approach implemented by SnapATAC (**Supplementary Fig. 8c**). Furthermore, the efficiency of the standard iterative LSI implementation in ArchR limits the requirement for this estimated LSI approach to only extremely large datasets (>200,000 cells for 32 GB RAM and 8 cores), whereas estimated LDM approaches are required for comparatively smaller datasets (>25,000 cells for 32 GB and 8 cores) in SnapATAC. ArchR therefore has the capacity to rapidly and efficiently analyze both large- and small-scale datasets.

### Robust inference of gene scores enables accurate cluster identification with ArchR

After clustering, investigators often aim to annotate the biological state related to each cluster. Methods for inferring gene expression from scATAC-seq data can generate “gene scores” of key marker genes that can enable accurate cluster annotation^5–8, 18^. However, the methods for integrating chromatin accessibility signal to generate these gene score predictions have not been extensively optimized. To this end, we used ArchR to benchmark 56 different models for inferring gene expression from scATAC-seq data using matched scATAC-seq and scRNA-seq data from PBMCs and bone marrow cells (**Fig. 2a and Supplementary Table 3**). To assess the performance of each model, we compared the known gene expression from previous methods integrating scATAC-seq with scRNA-seq^7, 11^ to the inferred gene scores derived from the model. By first establishing a rough linkage of ATAC-seq to RNA expression across many relatively diverse cell types (**Fig. 2a**), we could then determine which method for integrating ATAC-seq signal to predict gene expression had the best global performance across these data. The 56 gene score models varied by the regions included, the sizes of those regions, and the weights (based on genomic distance) applied to each region (**Fig. 2b and Supplementary Fig. 9a-h**). Models that incorporated ATAC-seq signal from the gene-body were more accurate than models that incorporated signal only from the promoter, likely due to the moderate increase in accessibility that occurs during active transcription. Moreover, incorporation of distal regulatory elements, weighted by distance, while accounting for the presence of neighboring genes (see methods) increased the accuracy of the gene score inference in all cases (**Supplementary Fig. 9a-h**). The most accurate model across both datasets was Model 42 (a model within the gene body extended + exponential decay + gene boundary class of models) (**Fig. 2b**) which integrates signal from the entire gene body, and scales signal with bi-directional exponential decays from the gene TSS (extended upstream by 5 kb) and the gene transcription termination site (TTS) while accounting for neighboring genes boundaries (**Fig. 2c**). This model yielded robust genome-wide gene score predictions in both PBMC and bone marrow cell datasets (**Fig. 2d-f and Supplementary Fig. 9i-j**). We additionally confirmed the efficacy of this class of gene score models using previously published paired bulk ATAC-seq and RNA-seq data from hematopoietic cells (**Supplementary Fig. 9k-m**)^19^. Given this analysis, we implemented this class of gene score models via Model 42 for all downstream analyses involving inferred gene expression in ArchR.

**Figure 2.**
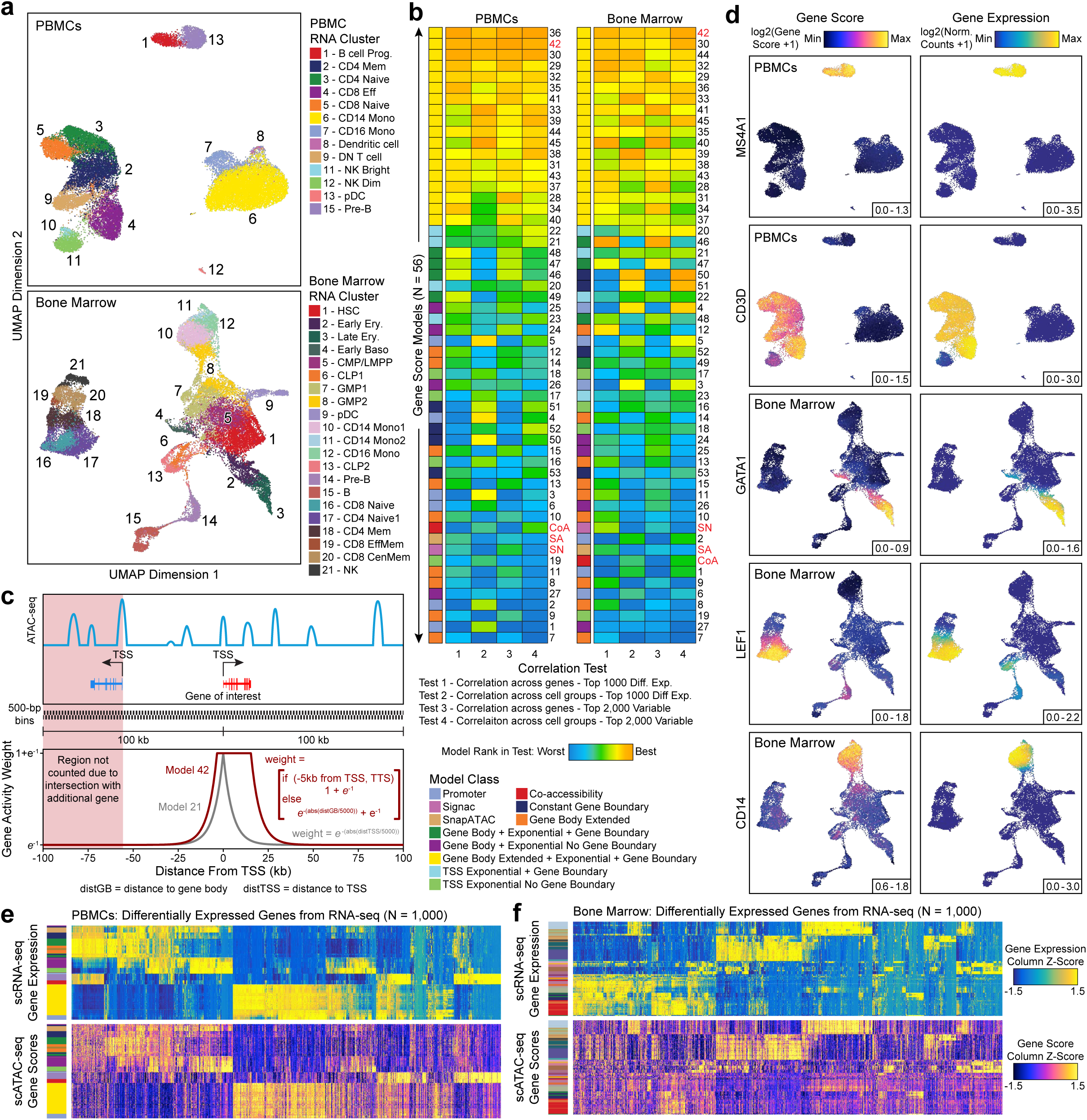
Optimized gene score inference models provide improved prediction of gene expression from scATAC-seq data. **a.** UMAPs of scATAC-seq data from (top) PBMCs and (bottom) bone marrow cells colored by aligned scRNA-seq clusters. This alignment is used for benchmarking all downstream scATAC-seq gene score models. **b.** Heatmaps summarizing the accuracy (measured by Pearson correlation) across 56 gene score models for both the top 1,000 differentially expressed and top 2,000 variable genes for both PBMC and bone marrow cell datasets. Each heatmap entry is colored by the model rank in the given correlation test as described below the heatmap. The model class is indicated to the left of each heatmap by color. SA, SnapATAC; SN, Signac; CoA, Co-accessibility. **c.** Illustration of the gene score Model 42, which uses bi-directional exponential decays from the gene TSS (extended upstream by 5 kb) and the gene transcription termination site (TTS) while accounting for neighboring gene boundaries (see methods). This model was shown to be more accurate than other models such as Model 21 which models an exponential decay from the gene TSS. **d.** Side-by-side UMAPs for PBMCs and bone marrow cells colored by (left) gene scores from Model 42 and (right) gene expression from scRNA-seq alignment for key immune cell-related marker genes. **e-f.** Heatmaps of (top) gene expression or (bottom) gene scores for the top 1,000 differentially expressed genes (selected from scRNA-seq) across all cell aggregates for (**e**) PBMCs or (**f**) bone marrow cells. Color bars to the left of each heatmap represent the PBMC or bone marrow cell cluster derived from scRNA-seq data.

### ArchR enables comprehensive analysis of massive-scale scATAC-seq data

ArchR is designed to handle datasets substantially larger (>1,000,000 cells) than those generated to date with modest computational resources. To illustrate this, we collected a compendium of high-quality published scATAC-seq data from immune cells generated with either the 10x Chromium system or the Fluidigm C1 system (49 samples, ∼220k cells; **Supplementary Figure 10a-d**). We refer to this compiled dataset as the hematopoiesis dataset. Using both a small-scale server infrastructure (8 cores, 32 GB RAM, with an HP Lustre file system) and a personal laptop (MacBook Pro laptop; 8 cores, 32 GB RAM, with an external USB hard drive), ArchR performed data import, dimensionality reduction, and clustering on ∼220k cells in less than three hours (**Fig. 3a and Supplementary Fig. 10e**). We next used ArchR to analyze a simulated set of over 1.2 million PBMCs, split into 200 individual samples. Under the same computational constraints, ArchR performed data import, dimensionality reduction, and clustering of more than 1.2 million cells in under 8 hours (**Fig. 3a and Supplementary Fig. 10e**).

**Figure 3.**
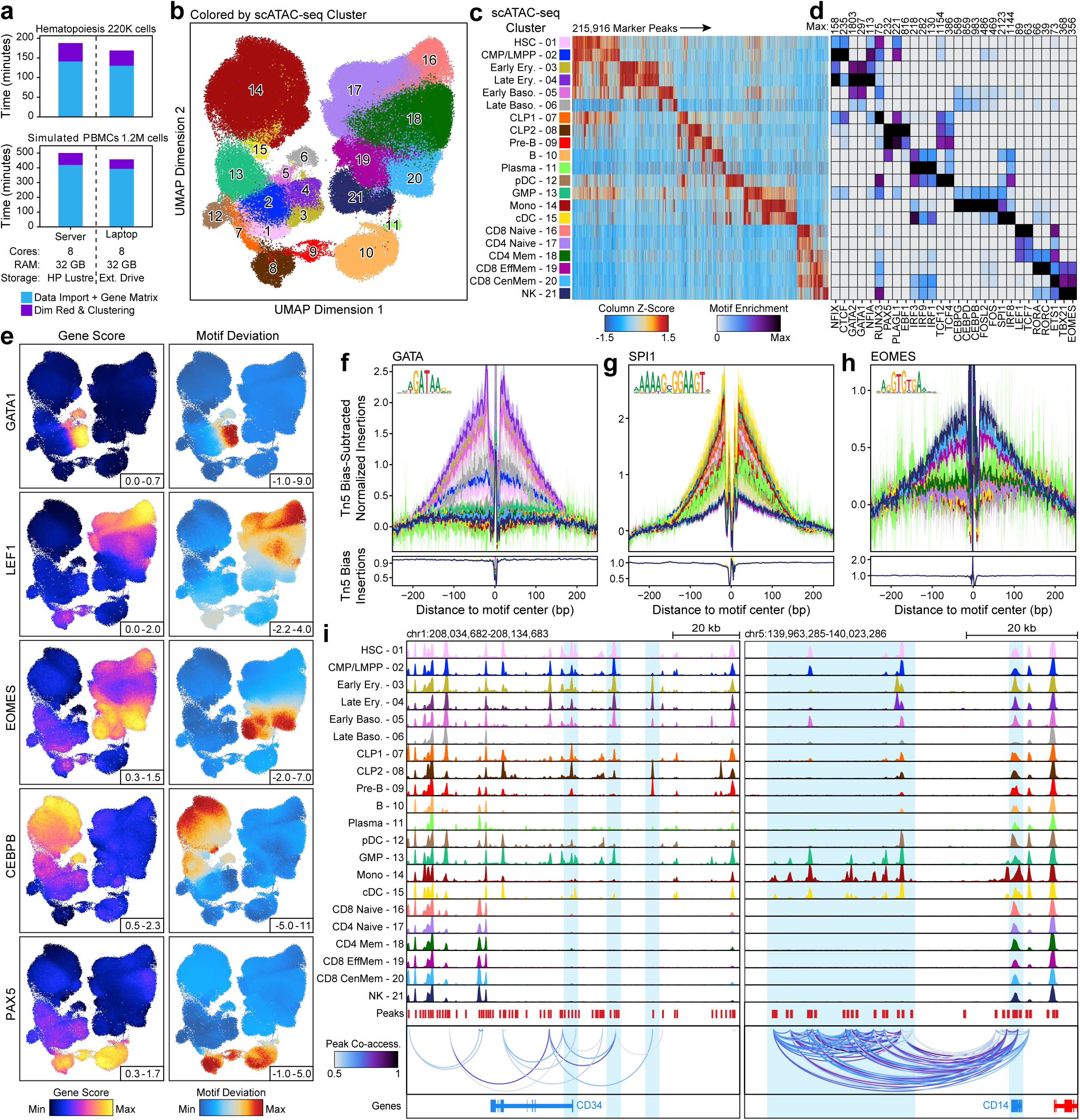
ArchR enables comprehensive analysis of massive-scale scATAC-seq data. **a.** Run times for ArchR-based analysis of over 220,000 and 1,200,000 single cells respectively using a small cluster-based computational environment (32 GB RAM and 8 cores with HP Lustre storage) and a personal MacBook Pro laptop (32 GB RAM and 8 cores with an external USB hard drive). Color indicates the relevant analytical step. **b.** UMAP of the hematopoiesis dataset colored by the 21 hematopoietic clusters. UMAP was constructed using LSI estimation with 25,000 landmark cells. **c.** Heatmap of 215,916 ATAC-seq marker peaks across all hematopoietic clusters identified with bias-matched differential testing. Color indicates the column Z-score of normalized accessibility. **d.** Heatmap of motif hypergeometric enrichment adjusted p-values within the marker peaks of each hematopoietic cluster. Color indicates the motif enrichment (-log10(p-value)) based on the hypergeometric test. **e.** Side-by-side UMAPs of (left) gene scores and (right) motif deviation scores for ArchR-identified TFs where the inferred gene expression is positively correlated with the chromVAR TF deviation across hematopoiesis. **f-h.** Tn5 bias-adjusted transcription factor footprints for GATA, SPI1, and EOMES motifs, representing positive TF regulators of hematopoiesis. Lines are colored by the 21 clusters shown in Figure 3c. **i.** Genome accessibility track visualization of marker genes with peak co-accessibility. (Left) *CD34* genome track (chr1:208,034,682-208,134,683) showing greater accessibility in earlier hematopoietic clusters (1-5, 7-8 and 12-13). (Right) *CD14* genome track (chr5:139,963,285-140,023,286) showing greater accessibility in earlier monocytic clusters (13-15).

Beyond these straightforward analyses, ArchR also provides an extensive suite of tools for more comprehensive analysis of scATAC-seq. Here we demonstrate these applications using the hematopoiesis dataset described above. Estimated LSI of this ∼220k-cell dataset recapitulated the overall structure of the data with a landmark dataset of as few as 500 cells (**Supplementary Fig. 10f**). Manual inspection of the resultant clusters with our uniform manifold approximation and projection (UMAP)^20^ led us to use the 25,000 cell landmark set (∼10% of total cells), which additionally showed minimal bias due to batch and data quality (**Fig. 3b and Supplementary Fig. 10g-i**). We identified 21 clusters spanning the hematopoietic hierarchy, calling clusters for even rare cell types such as plasma cells which comprise ∼0.1% (265 cells) of the total population. To generate a universal peak set from cluster-specific peaks, ArchR creates sample-aware pseudo-bulk replicates that recapitulate the biological variability within each cluster (**Supplementary Fig. 11a**). Peaks are then called from these pseudo-bulk replicates and a set of reproducible fixed-width non-overlapping peaks are identified using an iterative overlap merging procedure^21^ (**Supplementary Fig. 11b**). Using this approach, we identified 396,642 total reproducible peaks (**Supplementary Fig. 11c**), of which 215,916 are classified as differentially accessible peaks across the 21 clusters after bias-matched differential testing (see methods; **Fig. 3c**). Motif enrichment within these marker peaks revealed known TF regulators of hematopoiesis such as GATA1 in erythroid populations, CEBPB in monocytes, and PAX5 in B cell differentiation (**Fig. 3d**). In addition to motif enrichments, ArchR can calculate peak overlap enrichment with a compendium of previously published ATAC-seq datasets^19, 21–26^, identifying strong enrichment of peaks consistent with the cell type of each cluster (**Supplementary Fig. 11d**). To further characterize clusters, ArchR enables the projection of bulk ATAC-seq data into the single-cell-derived UMAP embedding^7^ via a down-sampling approach (**Supplementary Fig. 12a**). This allows for projection of sorted cell types, facilitating the identification of clusters based on well-validated bulk ATAC-seq profiles^19^ (**Supplementary Fig. 12b**). This projection analysis generates cell positions from bulk ATAC-seq data consistent with known cell types from a Fluidigm C1 scATAC-seq dataset of sorted hematopoietic cells including highly-similar hematopoietic stem and progenitor cells^4^ (**Supplementary Fig. 12c**) and aligns with inferred gene scores for canonical hematopoietic marker genes (**Supplementary Fig. 12d**).

ArchR also implements a scalable method for determination of transcription factor deviations from chromVAR^27^ in a sample independent manner (**Supplementary Fig. 12e**). TFs whose expression is highly correlated with their motif accessibility (i.e. putative positive regulators) can therefore be identified based on the correlation of the inferred gene score to the chromVAR motif deviation. This analysis identifies known drivers of hematopoietic differentiation such as GATA1 in erythroid populations, LEF1 in Naive T cell populations, and EOMES in NK/T Cell Memory populations. (**Fig. 3e, Supplementary Fig. 12f, and Supplementary Table 4**). ArchR also enables rapid footprinting of these TF regulators within clustered subsets while accounting for Tn5 biases^21^ using an improved C++ implementation (**Fig. 3f-h, Supplementary Fig. 12g-i**). Finally, ArchR identifies links between regulatory elements and target genes based on the co-accessibility of pairs of loci across single cells^1, 18^ (**Fig. 3i**).

### The interactive ArchR genome browser

In addition to these robust ATAC-seq analysis paradigms, ArchR provides a fully integrated and interactive genome browser (**Supplementary Fig. 13a**). The responsive and interactive nature of the browser is enabled by the optimized storage format within each Arrow file, providing support for dynamic cell grouping, track resolution, coloration, layout, and more. Launched via a single command, the ArchR browser enables cell cluster investigations of marker genes such as *CD34* for early hematopoietic stem and progenitor cells and *CD14* for monocytic populations (**Fig. 3i and Supplementary Fig. 13b-e**) while mitigating the need for external software for visualization of scATAC-seq data.

### ArchR enables integration of matched scRNA-seq and scATAC-seq datasets

ArchR also provides functionality to integrate scATAC-seq data with scRNA-seq data using Seurat’s infrastructure^11^. In brief, this integration requires matching the chromatin accessibility profiles and RNA expression for independent heterogeneous cells measured with two different assays. Single-cell epigenome-to-transcriptome integration is essential for understanding dynamic gene regulatory processes, and we anticipate this sort of analysis will become even more prevalent with the advent of platforms for simultaneous scATAC-seq and scRNA-seq. ArchR efficiently performs this cross-data alignment in parallel using slices of the scATAC-seq data (**Fig. 4a**). When performed on the hematopoiesis dataset, this integration enabled accurate scRNA-seq alignment for >220,000 cells in less than 1 hour (**Fig. 4b**). The alignment showed high concordance between linked gene expression and inferred gene scores for common hematopoietic marker genes (**Fig. 4c and Supplementary Fig. 14a**). Using this cross-platform alignment, ArchR also provides methods to identify putative cis-regulatory elements based on correlated peak accessibility and gene expression^7, 21^ (**Supplementary Fig. 15a**). In the example hematopoiesis dataset, this analysis identified 70,239 significant peak-to-gene linkages across the hematopoietic hierarchy (**Supplementary Fig. 15b and Supplementary Table 5**).

**Figure 4.**
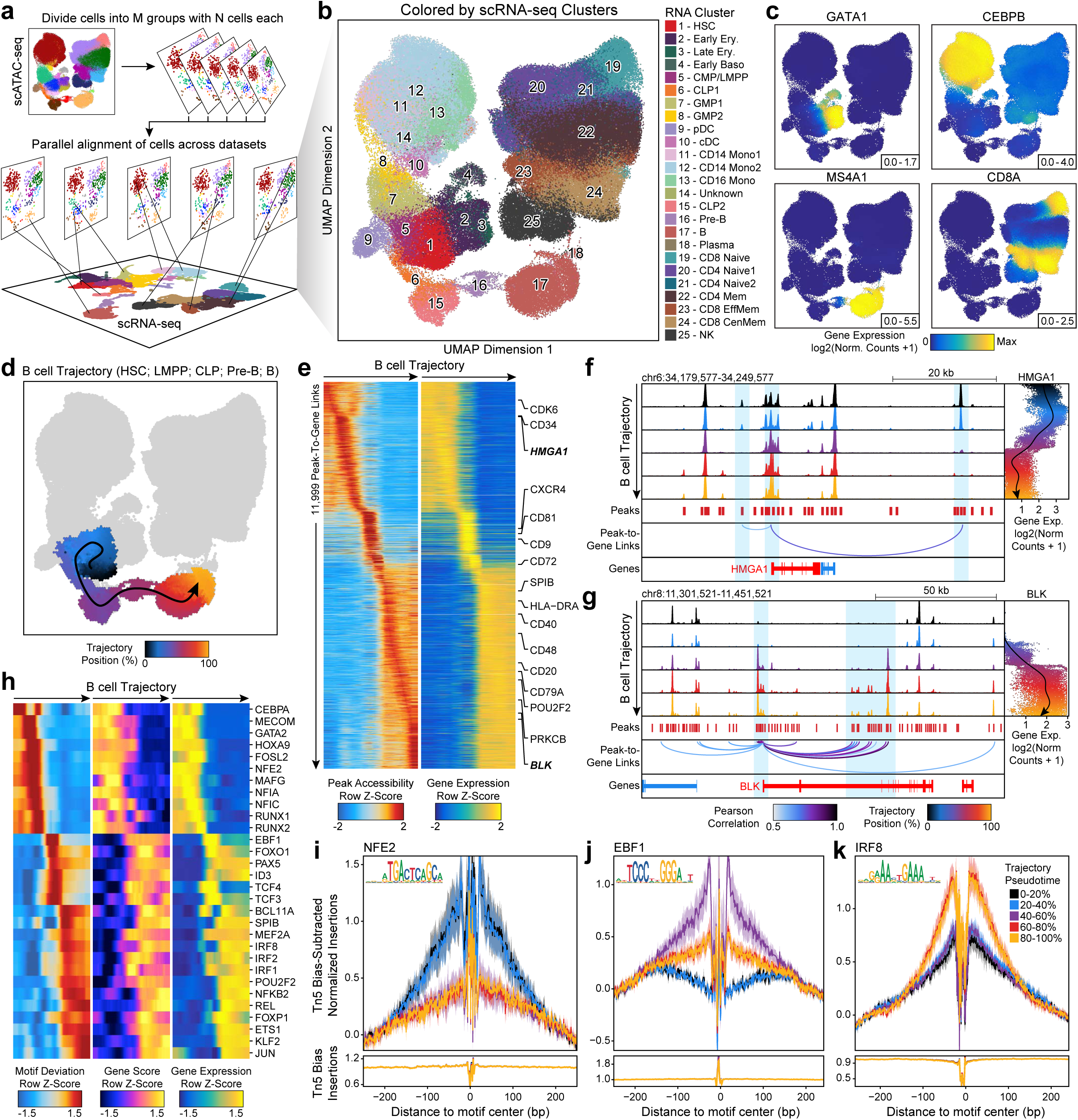
Integration of scATAC-seq and scRNA-seq data by ArchR identifies gene regulatory trajectories of hematopoietic differentiation. **a.** Schematic of scATAC-seq alignment with scRNA-seq data in M slices of N single cells. These slices are independently aligned to a reference scRNA-seq dataset and then the results are combined for downstream analysis. This integrative design facilitates rapid large-scale integration with low-memory requirements. **b-d.** UMAP of scATAC-seq data from the hematopoiesis dataset colored by (**b**) alignment to previously published hematopoietic scRNA-seq-derived clusters, (**c**) integrated scRNA-seq gene expression for key marker TFs and genes, or (**d**) cell alignment to the ArchR-defined B cell trajectory. In (**d**), the smoothed arrow represents a visualization of the interpreted trajectory (determined in the LSI subspace) in the UMAP embedding. **e.** Heatmap of 11,999 peak-to-gene links identified across the B cell trajectory with ArchR. **f-g.** Genome track visualization of the (**f**) *HMGA1* locus (chr6:34,179,577-34,249,577) and (**g**) *BLK* locus (chr8:11,301,521-11,451,521). Single-cell gene expression across pseudo-time in the B cell trajectory is shown to the right. Inferred peak-to-gene links for distal regulatory elements across the hematopoiesis dataset is shown below. **h.** Heatmap of positive TF regulators whose gene expression is positively correlated with chromVAR TF deviation across the B cell trajectory. **i-k.** Tn5 bias-adjusted transcription factor footprints for (**i**) NFE2, (**j**) EBF1, and (**k**) IRF8 motifs, representing positive TF regulators across the B cell trajectory. Lines are colored by the position in pseudo-time of B cell differentiation.

Finally, ArchR facilitates cellular trajectory analysis to identify the predicted path of gene regulatory changes from one set of cells to another, a unique type of insight enabled by single-cell data. To carry out this analysis, ArchR initially creates a cellular trajectory based on a sequence of user-supplied clusters or groups. ArchR then identifies individual cell positions along this trajectory based on Euclidean distance within an N-dimensional subspace^6^. Using B cells as an example, ArchR traces cells along the B cell differentiation trajectory and identifies 11,999 peak-to-gene links that have correlated regulatory dynamics across the B cell differentiation (**Fig. 4e**). Sequencing tracks of the *HMGA1* locus, active in stem and progenitor cells, and the *BLK* locus, active in differentiated B cells, demonstrate how accessibility at linked peaks correlates with longitudinal changes in gene expression across pseudo-time (**Fig. 4f-g**). Moreover, using this same paradigm, ArchR can identify TF motifs with accessibility that are positively correlated with gene expression of TF genes across the same B cell trajectory (**Fig. 4h**). Transcription factor footprinting of a subset of these TFs further illustrates the dynamics in the local accessibility at the binding sites of these lineage-defining TFs across B cell differentiation pseudo-time (**Fig. 4i-k**).

## DISCUSSION

Chromatin accessibility data provides a lens through which we can observe the gene regulatory programs that underlie cellular state and identity. The highly cell type-specific nature of cis-regulatory elements makes profiling of single-cell chromatin accessibility an attractive method to understand cellular heterogeneity and the molecular processes underlying complex control of gene expression. With the advent of methods to profile chromatin accessibility across thousands of single cells, scATAC-seq has quickly become a method-of-choice for many single-cell applications. However, compared to scRNA-seq, analysis of scATAC-seq data remains comparatively immature with no clear standards, thus dissuading many from adopting this informative technique.

To address this unmet need, we developed ArchR, an end-to-end software solution that will greatly expedite single-cell chromatin analysis for any biologist. Low memory usage, parallelized operations, and an intuitive and user-focused, yet extensive and powerful tool suite make ArchR an ideal platform for scATAC-seq data analysis. In contrast to currently available software packages, ArchR is designed to handle millions of cells using commonly available computational resources, such as a laptop running a Unix-based operating system. As such, ArchR provides the analytical support necessary for the massive scale of ongoing efforts to catalog the compendium of diverse cell types throughout the body at single-cell resolution^28^. In addition to the dramatic improvements in run time, memory efficiency, and scale, ArchR supports state-of-the-art chromatin-based analyses including genome-wide inference of gene activity, transcription factor footprinting, and data integration with matched scRNA-seq, enabling statistical linkage of cis- and trans-acting regulatory factors to gene expression profiles. Moreover, the performance improvements from ArchR enable interactive data analysis whereby end-users can iteratively adjust analytical parameters and thus optimize identification of biologically meaningful results. This is especially important in the context of single-cell data where a one-size-fits-all analytical pipeline is not relevant or desirable. Supervised identification of clusters, resolution of subtle batch effects, and biology-driven data exploration are intrinsically necessary for a successful scATAC-seq analysis and ArchR supports these efforts by enabling rapid analytical processes. ArchR provides an open-source analysis platform with the flexibility, speed, and power to support the rapidly increasing efforts to understand complex tissues, organisms, and ecosystems at the resolution of individual cells.

## Methods

### Code Availability and Documentation

Extensive documentation and a full user manual are available at www.ArchRProject.com. The software is open-source and all code can be found on GitHub at https://github.com/GreenleafLab/ArchR. Additionally, code for producing the majority of analyses from this paper is available at the publication page https://github.com/GreenleafLab/ArchR_2020.

### Data Availability

Bulk and scATAC-seq data from the cell line mixing experiment will be available through GEO (accession number in progress). All other scATAC-seq data used were from publicly available sources as outline in **Supplementary Table 1**. We additionally have made available other analysis files on our publication page https://github.com/GreenleafLab/ArchR_2020.

### Genome and Transcriptome Annotations

All analyses were performed with the hg19 genome (except the Mouse Atlas with mm9). R-based analysis used the BSgenome package with “BSgenome.Hsapiens.UCSC.hg19” (“BSgenome.Mmusculus.UCSC.mm9” for Mouse Atlas) for genomic coordinates and the TxDb package with “TxDb.Hsapiens.UCSC.hg19.knownGene” (“TxDb.Mmusculus.UCSC.mm9.knownGene” for Mouse Atlas) gene annotations unless otherwise stated.

### Cell Type Abbreviations

In many of the figure legends, abbreviations are used for cell types of the hematopoietic system. HSC, hematopoietic stem cell; LMPP, lymphoid-primed multipotent progenitor cell; CMP, common myeloid progenitor; CLP, common lymphoid progenitor; GMP, granulocyte macrophage progenitor; CD4 Mem, CD4 memory T cell; CD4 Naive, CD4 naïve T cell; CD8 Naive, CD8 naïve T cell; CD8 Eff, CD8 effector T cell; CD8 EffMem, CD8 effector memory T cell; CD8 CenMem, CD8 central memory T cell; Mono, monocyte; pDC, plasmacytoid dendritic cell; NK, natural killer cell; Ery, erythroid; Baso, basophil.

### scATAC-seq Data Generation – Cell Lines

With the exception of MCF10A, all cell lines were cultured in the designated media containing 10% FBS and penicillin/streptomycin. Jurkat, THP1, and K562 cell lines were ordered from ATCC and cultured in RPMI-1640. GM12878 cells were ordered from Coriell and cultured in RPMI-1640. HeLa, HEK-293T, and HT1080 cell lines were ordered from ATCC and cultured in DMEM. T24 cells were ordered from ATCC and cultured in McCoy’s 5A. MCF7 cells were ordered from ATCC and cultured in EMEM containing 0.01 mg/ml of human insulin (Millipore-Sigma 91077C). MCF10A cells were ordered from ATCC and cultured in DMEM/F12 containing 5% horse serum (Thermo Fisher 16050130), 0.02 ug/ml human EGF (PeproTech AF-100-15), 0.5 ug/ml hydrocortisone (Millipore-Sigma H0888), 0.1 ug/ml Cholera toxin (Millipore-Sigma C8052), 10 ug/ml insulin from bovine pancreas (Millipore-Sigma I6634), and penicillin/streptomycin. Cultured cells were viably cryopreserved in aliquots of 100,000 cells using 100 ul of BAMBANKER freezing media (Wako Chemicals 302-14681) so that scATAC-seq could be performed on all cells at the same time. For each cell line, cells were thawed via the addition of 1 mL ice-cold resuspension buffer (RSB) [10 mM Trish-HCl pH 7.4, 10 mM NaCl, 3 mM MgCl2] containing 0.1% Tween-20 (RSB-T). Cells were pelleted in a fixed-angle rotor at 300 RCF for 5 minutes at 4°C. The supernatant was removed and the pellet was resuspended in 100 uL of ice-cold lysis buffer (RSB containing 0.1% Tween-20, 0.1% NP-40, and 0.01% digitonin) and incubated on ice for 3 minutes. To dilute lysis, 1 mL of chilled RSB-T was added to each tube and the cells were pelleted as before. The supernatant was removed and the pelleted nuclei were resuspended in Diluted Nuclei Buffer (10x Genomics). The nuclei stock concentration was determined for each cell line using Trypan Blue and a total of 5,000 nuclei from each cell line were pooled together and loaded into the 10x Genomics scATAC-seq (v1) transposition reaction. The remainder of the scATAC-seq library preparation was performed as recommended by the manufacturer. Resultant libraries were sequenced on an Illumina NovaSeq6000 using an S4 flow cell and paired-end 99-bp reads. In addition to this pooled scATAC-seq, each cell line was used to generate bulk ATAC-seq libraries as described previously^26^. Bulk ATAC-seq libraries were pooled and purified via PAGE gel prior to sequencing on an Illumina HiSeq4000 using paired-end 75-bp reads.

### scATAC-seq Processing – Cell Line Mixing

Raw sequencing data was converted to FASTQ format using cellranger-atac mkfastq (10x Genomics, version 1.0.0). Single-cell ATAC-seq reads were aligned to the hg19 reference genome (https://support.10xgenomics.com/single-cell-atac/software/downloads/latest) and quantified using cellranger-count (10x Genomics, version 1.0.0). Genotypes used to perform demuxlet were determined as follows for each cell line: Bulk ATAC-seq FASTQ files were processed and aligned using PEPATAC (http://code.databio.org/PEPATAC/) as described previously^21^. Peaks were identified using MACS2 and a union set of variable-width accessible regions was identified using bedtools merge (version 2.26.0). These accessible regions were genotyped across all samples using samtools mpileup (version 1.5) and Varscan mpileup2snp (version 2.4.3) with the following parameters “--min-coverage 5 --min-reads2 2 --min-var-freq 0.1 --strand-filter 1 --output-vcf 1”. All positions containing a single nucleotide variant were compiled into a master set and then each cell line was genotyped at those specific single-base locations using samtools mpileup. The allelic depth at each position was converted into a quaternary genotype (homozygous A, heterozygous AB, homozygous B, or insufficient data to generate a confident call). Then, for each cell line, inferred genotype probabilities were created based on those quaternary genotypes and a VCF file was created for input to demuxlet using recommended parameters. Demuxlet was used to identify the cell line of origin for individual cells and to identify doublets based on mixed genotypes.

### ArchR Methods – Preface

All ArchR features were carefully designed and optimized to enable analysis of 250,000 cells or greater on a minimal computing environment in R. All ArchR HDF5-formatted processing was performed with the Bioconductor^29^ package “rhdf5” (https://www.bioconductor.org/packages/release/bioc/html/rhdf5.html). All ArchR genomic coordinate operations were performed with the Bioconductor package “GenomicRanges” (https://bioconductor.org/packages/release/bioc/html/GenomicRanges.html) and “IRanges” (https://bioconductor.org/packages/release/bioc/html/IRanges.html).

### ArchR Methods – scATAC Definitions

**Fragments –** In ATAC-seq data analysis, a “fragment” refers to a sequenceable DNA molecule created by two transposition events. Each end of that fragment is sequenced using paired-end sequencing. The inferred single-base position of the start and end of the fragment is adjusted based on the insertion offset of Tn5. As reported previously^30^, Tn5 transposase binds to DNA as a homodimer with 9-bp of DNA between the two Tn5 molecules. Because of this interaction, each Tn5 homodimer binding event creates two insertions, separated by 9 bp. Thus, the actual central point of the “accessible” site is in the very center of the Tn5 dimer, not the location of each Tn5 insertion. To account for this, we apply an offset to the individual Tn5 insertions, adjusting plus-stranded insertion events by +4 bp and minus-stranded insertion events by -5 bp. This offset is consistent with the convention put forth during the original description of ATAC-seq^31^. Thus, in ArchR, “fragments” refers to a table or Genomic Ranges object containing the chromosome, offset-adjusted chromosome start position, offset-adjusted chromosome end position, and unique cellular barcode ID corresponding to each sequenced fragment.

**Tn5 insertions –** In ArchR, “insertions” refers to the offset-adjusted single-base position of Tn5 insertion on either end of the fragment. Insertion positions are accessed in ArchR primarily using resize(fragments, 1, “start”) and resize(fragments, 1, “end”). See the description of “fragments” above for a detailed description of Tn5 insertion offsets.

**Counting Accessibility –** In ArchR, “counting accessibility” refers to counting the number of Tn5 insertions observed within each described feature.

**TSS enrichment score –** In ArchR, the “TSS enrichment” refers to the relative enrichment of Tn5 insertions at gene TSS sites genome-wide compared to a local background. This represents a measure of signal-to-background in ATAC-seq data. See below for how TSS enrichment is calculated in ArchR. In this work, the TSS enrichment score from ArchR is based on the TSS regions defined by the TxDb.Hsapiens.UCSC.hg19.knownGene (or TxDb.Mmusculus.UCSC.mm9.knownGene for the Mouse Atlas) transcript database object.

### ArchR Methods – Arrow Files and ArchRProject

The base unit of an analytical project in ArchR is called an “Arrow file”. Each Arrow file stores all of the data associated with an individual sample (i.e. metadata, accessible fragments, and data matrices). Here, an “individual sample” would be the most detailed unit of analysis desired (for ex. a single replicate of a particular condition). During creation and as additional analyses are performed, ArchR updates and edits each Arrow file to contain additional layers of information. It is worth noting that, to ArchR, an Arrow file is actually just a path to an external file stored on disk. More explicitly, an Arrow file is not an R-language object that is stored in memory but rather an HDF5-format file stored on disk. Because of this, we use an “ArchRProject” object to associate these Arrow files together into a single analytical framework that can be rapidly accessed in R. This ArchRProject object is small in size and is stored in memory.

Certain actions can be taken directly on Arrow files while other actions are taken on an ArchRProject which in turn updates each associated Arrow file. Because Arrow files are stored as large HDF5-format files, “get-er” functions in ArchR retrieve data by interacting with the ArchRProject while “add-er” functions either (i) add data directly to Arrow files, (ii) add data directly to an ArchRProject, or (iii) add data to Arrow files by interacting with an ArchRProject.

### ArchR Methods – Reading Input Data into an Arrow File

ArchR can utilize multiple input formats of scATAC-seq data which is most frequently in the format of fragment files and BAM files. Fragment files are tabix-sorted text files containing each scATAC-seq fragment and the corresponding cell ID, one fragment per line. BAM files are binarized tabix-sorted files that contain each scATAC-seq fragment, raw sequence, cellular barcode id and other information. The input format used will depend on the pre-processing pipeline used. For example, the 10x Genomics Cell Ranger software returns fragment files while sci-ATAC-seq applications would use BAM files. Given a specified genome annotation (ArchR has pre-loaded genome annotations for mm9, mm10, hg19, and hg38 and additional genomes can be added manually), ArchR reads these input files in sub-chromosomal chunks using Rsamtools. ArchR uses “scanTabix” to read fragment files and “scanBam” to read BAM files. During this input process, each input chunk is converted into a compressed table-based representation of fragments containing each fragment chromosome, offset-adjusted chromosome start position, offset-adjusted chromosome end position and cellular barcode ID. These chunk-wise fragments are then stored in a temporary HDF5-formatted file to preserve memory usage while maintaining rapid access to each chunk. Finally, all chunks associated with each chromosome are read, organized, and re-written to an “Arrow file” within a single HDF5 group called “fragments”. This pre-chunking procedure enables ArchR to process extremely large input files efficiently and with low memory usage, enabling full utilization of parallel processing.

### ArchR Methods – QC Based on TSS Enrichment and Unique Nuclear Fragments

Strict quality control (QC) of scATAC-seq data is essential to remove the contribution of low-quality cells. In ArchR, one characteristic of “low-quality” is a low signal-to-background ratio, which is often attributed to dead or dying cells which have de-chromatinzed DNA which allows for random transposition genome-wide. Traditional bulk ATAC-seq analysis has used the TSS enrichment score as part of a standard workflow (https://www.encodeproject.org/atac-seq/) for determination of signal-to-background. We and others have found the TSS enrichment to be representative across the majority of cell types tested in both bulk ATAC-seq and scATAC-seq. The idea behind the TSS enrichment score metric is that ATAC-seq data is universally enriched at gene TSS regions compared to other genomic regions. By looking at per-base-pair accessibility centered at these TSS regions, we see a local enrichment relative to flanking regions (1900-2000 bp distal in both directions). The ratio between the peak of this enrichment (centered at the TSS) relative to these flanking regions represents the TSS enrichment score. Traditionally, the per-base-pair accessibility is computed for each bulk ATAC-seq sample and then this profile is used to determine the TSS enrichment score. Performing this operation on a per-cell basis in scATAC-seq is relatively slow and computationally expensive. To accurately approximate the TSS enrichment score per single cell, we count the average accessibility within a 50-bp region centered at each single-base TSS position and divide this by the average accessibility of the TSS flanking positions (+/- 1900 – 2000 bp). This approximation was highly correlated (R > 0.99) with the original method and values were extremely close in magnitude. By default in ArchR, pass-filter cells are identified as those cells having a TSS enrichment score greater than 4 and more than 1000 unique nuclear fragments (i.e those fragments that do not map to chrM).

### ArchR Methods – Tile Matrix

Traditional bulk ATAC-seq analysis relies on the creation of a peak matrix from a peak-set encompassing the precise accessible regions across all samples. This peak set, and thus the resulting peak matrix, is specific to the samples used in the analysis and must be re-generated when new samples are added. Moreover, identification of peaks from scATAC-seq data would optimally be performed after clusters were identified to ensure that cluster-specific peaks are captured. Thus, the optimal solution for scATAC-seq would be to identify an unbiased and consistent way to perform analysis prior to cluster identification, without the need for calling peaks. xbecause this bin size approximates the size of most regulatory elements. To circumvent the requirement for calling peaks prior to cluster identification, others have tiled the genome into fixed non-overlapping tiled windows. This method additionally benefits from being stable across samples and the tiled regions do not change based on inclusion of additional samples. However, these tiled windows are usually greater than or equal to 5 kb in length, which is more than 10-fold greater than the size of typical accessible regions containing TF binding sites^15–17^. For this reason, ArchR uses 500-bp genome-wide tiled windows for all analysis upstream of cluster identification. To create a tile matrix, ArchR reads in the scATAC-seq fragments for a chromosome and converts these to insertions. ArchR then floors these insertions to the nearest tile region with floor(insertion / tileSize) + 1. The tile regions and cell barcode id (as an integer) are then used as input for Matrix::sparseMatrix which tallies the number of input rows (tiles, denoted as i) and columns (cells, denoted as j) and creates a sparseMatrix. This analysis is performed for each chromosome and stored in the corresponding Arrow file. This fast and efficient conversion of scATAC-seq fragments to a tile matrix, without computing genomic overlaps, facilitates efficient construction of 500-bp tile matrices for analyses.

### ArchR Methods – Gene Score Matrix

ArchR facilitates the inference of gene expression from chromatin accessibility (called “gene scores”) by using custom distance-weighted accessibility models. For each chromosome, ArchR creates a tile matrix (user-defined tile size that is not pre-computed, default is 500 bp), overlaps these tiles with the gene window (user-defined, default is 100 kb), and then computes the distance from each tile (start or end) to the gene body (with optional extensions upstream or downstream) or gene start. We have found that the best predictor of gene expression is the local accessibility of the gene region which includes the promoter and gene body (**Supplementary Fig. 9**). To properly account for distal accessibility, for each gene ArchR identifies the subset of tiles that are within the gene window and do not cross another gene region. This filtering allows for inclusion of distal regulatory elements that could improve the accuracy of predicting gene expression values but excludes regulatory elements more likely to be associated with another gene (for ex. the promoter of a nearby gene). The distance from each tile to the gene is then converted to a distance weight using a user-defined accessibility model (default is *e*^(-abs(distance)/5000)^ + *e*^-1^). When the gene body is included in the gene region (where the distance-based weight is the maximum weight possible), we found that extremely large genes can bias the overall gene scores. In these cases, the total gene scores can vary substantially due to the inclusion of insertions in both introns and exons. To help adjust for these large differences in gene size, ArchR applies a separate weight for the inverse of the gene size (1 / gene size) and scales this inverse weight linearly from 1 to a hard max (which can be user-defined, with a default of 5). Smaller genes thus receive larger relative weights, partially normalizing this length effect. The corresponding distance and gene size weights are then multiplied by the number of Tn5 insertions within each tile and summed across all tiles within the gene window (while still accounting for nearby gene regions as described above). This summed accessibility is a “gene score” and is depth normalized across all genes to a constant (user-defined, default of 10,000). Computed gene scores are then stored in the corresponding Arrow file for downstream analyses.

### ArchR Methods – Iterative LSI Procedure

The default LSI implementation in ArchR is conceptually similar to the method introduced in Signac (https://satijalab.org/signac/), which, for a cell x features matrix (typically tiles or peaks), uses a term frequency (column sums) that has been depth normalized to a constant (10,000) followed by normalization with the inverse document frequency (1 / row sums) and then log-transformed (aka log(TF-IDF)). This normalized matrix is then factorized by singular value decomposition (SVD) and then standardized across the reduced dimensions for each cell via z-score. ArchR additionally allows for the use of alternative LSI implementations based on previously published scATAC-seq papers^5–7^. As mentioned above, the input to LSI-based dimensionality reduction is the genome-wide 500-bp tile matrix.

In scRNA-seq, identifying variable genes is a common way to compute dimensionality reduction (such as PCA), as these highly variable genes are more likely to be biologically important, and focusing on these genes likely reduces low-level contributions of variance potentially due to experimental noise. ScATAC-seq data is binary, precluding the possibility of identifying variable peaks for dimensionality reduction. Therefore, rather than identifying the most variable peaks, we initially tried using the most accessible features as input to LSI; however, the results when running multiple samples exhibited a high degree of noise and low reproducibility. We therefore moved to our previously described “iterative LSI” approach^6, 7^. This approach computes an initial LSI transformation on the most accessible tiles and identifies lower resolution clusters that are driven by clear biological differences. For example, when performed on peripheral blood mononuclear cells, this approach will identify clusters corresponding to the major cell types (T cells, B cells, and monocytes). Then ArchR computes the average accessibility for each of these clusters across all features creating “pseudo-bulks”. ArchR then identifies the most variable peaks across these pseudo-bulks to use as features for the second round of LSI. In this second iteration, the selected variable peaks correspond more similarly to the variable genes used in scRNA-seq LSI implementations, insofar as they are highly variable across biologically meaningful clusters. We have found this approach can also effectively minimize batch effects and allows operations on a more reasonably sized feature matrix. Additionally, we observe that this procedure still allows the identification of rare cell types, such as plasma cells in the bone marrow cell dataset that exist at ∼0.1% prevalence. For larger batch effects, ArchR enables Harmony-based batch correction on the LSI-reduced coordinates^32^.

### ArchR Methods – Estimated LSI Procedure

For extremely large scATAC-seq datasets, ArchR can estimate the LSI dimensionality reduction with LSI projection. This procedure is similar to the iterative LSI workflow, however the LSI procedure differs. First, a subset of randomly selected “landmark” cells is used for LSI dimensionality reduction. Second, the remaining cells are TF-IDF normalized using the inverse document frequency determined from the landmark cells. Third, these normalized cells are projected into the SVD subspace defined by the landmark cells. This leads to an LSI transformation based on a small set of cells used as landmarks for the projection of the remaining cells. This estimated LSI procedure is efficient with ArchR because, when projecting the new cells into the landmark cells LSI, ArchR iteratively reads in the cells from each sample and LSI projects them without storing them all in memory. This optimization leads to minimal memory usage and further increases the scalability for extremely large datasets. Even with comparatively small landmark cell subsets (500-5000 cells), we find that this procedure is able to maintain the global structure and recapitulates the clusters well; however, the required landmark set size is dependent on the proportion of different cells within the dataset.

### ArchR Methods – Identification of Doublets

Single-cell data generated on essentially any platform is susceptible to the presence of doublets. A doublet refers to a single nano-reaction (i.e. a droplet) that received a single barcoded bead and more than one cell/nucleus. This causes the reads from more than one cell to appear as a single cell. For 10x Genomics applications, the percentage of total “cells” that are actually doublets is proportional to the number of cells loaded into the reaction. Even at lower cell loadings as recommended by standard kit use, more than 5% of the data may come from doublets, and this spurious data exerts substantial effects on clustering. This issue becomes particularly problematic in the context of developmental/trajectory data because doublets can look like a mixture between two cell types and this can be confounded with intermediate cell types or cell states.

To predict which “cells” are actually doublets in ArchR, we synthesize in silico doublets from the data by mixing the reads from thousands of combinations of individual cells. Next, we perform iterative LSI followed by UMAP for each individual sample. We then LSI project the synthetic doublets into the LSI subspace followed by UMAP projection. ArchR identifies the *k*-nearest neighbors (user-defined, default 10) to each simulated projected doublet. By iterating this procedure N times (user-defined, default 3 times the total number of cells), we can compute binomial enrichment statistics (assuming every cell could be a doublet with equal probability) for each single cell based on the presence of nearby simulated projected doublets (in the LSI or UMAP subspace defined by the user). This approach is similar to previous approaches^12, 13^, but differs in that LSI is used for dimensionality reduction and UMAP projection is used for identification. The number of doublets to remove is then determined based on either the number of cells that pass QC or for the approximate number of cells loaded as defined by the user. While we have optimized these parameters for general use, users should sensibly check their results with and without doublet removal.

### ArchR Methods – Identification of Clusters

ArchR uses established scRNA-seq clustering methods that use graph clustering on the LSI dimensionality reduction coordinates to resolve clusters. By default, ArchR uses Seurat’s graph clustering with “Seurat::FindClusters” for identifying high fidelity clusters^11^. ArchR additionally supports scran^33^ for single-cell clustering.

### ArchR Methods – t-SNE and UMAP Embeddings

ArchR supports both t-distributed stochastic neighbor embedding (t-SNE) and uniform manifold approximation and projection (UMAP) single-cell embedding methodologies. ArchR uses previously determined reduced dimensions as input for these embeddings. t-SNE analysis is performed using the “Rtsne” package in R by default. UMAP analysis is performed using the “uwot” package in R by default. The results are stored within an ArchRProject and then used for plotting and subsequent analyses.

### ArchR Methods – Sample-Aware Pseudo-Bulk Replicate Generation

Because of the sparsity of scATAC-seq data, operations are often performed on aggregated groups of single cells. Most frequently, these groups are defined by clustering, and it is assumed that each local cluster represents a relatively homogeneous cell type or cell state. This process of combining data from multiple individual cells creates “pseudo-bulk” data, because it resembles the data derived from a bulk ATAC-seq experiment.

A feature unique to ArchR is the creation of sample-aware pseudo-bulk replicates from each cell group to use for performing statistical tests (such as reproducible peak identification or TF footprinting). ArchR does this via a complex decision tree which is dependent upon a user-specified desired number of replicates and number of cells per replicate as presented in **Supplementary Fig. 11a**. Briefly, ArchR attempts to create pseudo-bulk replicates in a sample-aware fashion. This means that each individual pseudo-bulk replicate only contains cells from a single biological sample. This feature enables the preservation of variability associated with biological replicates. If the desired number of replicates cannot be created in this fashion, ArchR uses progressively less stringent requirements to create the pseudo-bulk replicates. First, ArchR attempts to create as many pseudo-bulk replicates in a sample-aware fashion as possible and then create the remaining pseudo-bulk replicates in a sample-agnostic fashion by sampling without replacement. If this is not possible, ArchR attempts to create the desired number of pseudo-bulk replicates in a sample-agnostic fashion by sample without replacement across all samples. If this is not possible, ArchR attempts the same procedure by sampling without replacement within a single replicate but with replacement across different replicates without exceeding a user-specified sampling ratio. If all of these attempts fail, ArchR will create the specified number of pseudo-bulk replicates by sampling with replacement within a single replicate and with replacement across different replicates. The fragments from all cells within a pseudo-bulk replicate are converted to insertions and to a run-length encoding (RLE) coverage object using the “coverage” function in R. This insertion coverage object (similar to a bigwig) is then written to a separate HDF5-formatted coverage file. ArchR next identifies single-base resolution Tn5 insertion sites for each pseudo-bulk replicate, resizes these 1-bp sites to k-bp (user-defined, default is 6) windows (-k/2 and + (k/2 - 1) bp from insertion), and then creates a k-mer frequency table using the “oligonucleotidefrequency(w=k, simplify.as=”collapse”)” function from the Biostrings package. ArchR then calculates the expected k-mers genome-wide using the same function with the BSGenome-associated genome file. These Tn5 k-mer values represent the Tn5 bias genome-wide and are then stored in the pseudo-bulk replicate HDF5 coverage file. This coverage file contains similar information to a bigwig file with Tn5 insertion bias but in a fast-access HDF5 format. This coverage file can be used for peak-calling and TF footprinting with Tn5 bias correction.

### ArchR Methods – Peak Calling

In ArchR, peak calling is performed on the HDF5-format pseudo-bulk-derived coverage files described above. By default, ArchR calls peak summits with MACS2 using single-base insertion positions derived from the coverage files (written to a bed file with data.table) with user-specified values for MACS2 parameters including gsize, shift (default -75), and extsize (default 150) along with the “nomodel” and “nolambda” flags. These single-base peak summit locations are extended to a 501-bp width. We use 501-bp fixed-width peaks because they make downstream computation easier as peak length does not need to be normalized. Moreover, the vast majority of peaks in ATAC-seq are less than 501-bp wide. Using variable-width peaks also makes it difficult to merge peak calls from multiple samples without creating extremely large peaks that create confounding biases.

To create a merged non-overlapping fixed-width union peak set, ArchR implements an iterative overlap removal procedure that we introduced previously^21^. Briefly, peaks are first ranked by their significance, then the most significant peak is retained and any peak that directly overlaps with the most significant peak is removed from further analysis. This process is repeated with the remaining peaks until no more peaks exist. This procedure avoids daisy-chaining and still allows for use of fixed-width peaks. We use a normalized metric for determining the significance of peaks because the reported MACS2 significance is proportional to the sequencing depth. This process is outlined in **Supplementary Fig. 11b**.

### ArchR Methods – Interactive Genome Browser

One challenge inherent to scATAC-seq data analysis is genome-track level visualizations of chromatin accessibility observed within groups of. Traditionally, track visualization requires grouping the scATAC-seq fragments, creating a genome coverage bigwig, and normalizing this track for quantitative visualization. Typically, end-users use a genome browser such as the WashU Epigenome Browser, the UCSC Genome Browser, or the IGV browser to visualize these sequencing tracks. This process involves using multiple software and any change to the cellular groups or addition of more samples requires re-generation of bigwig files etc., which can become time consuming. For this reason, ArchR has a Shiny-based interactive genome browser that can be launched with a simple line of code “ArchRBrowser(ArchRProj)”. The data storage strategy implemented in Arrow files allows this interactive browser to dynamically change the cell groupings, resolution, and normalization, enabling real-time track-level visualizations. The ArchR Genome Browser also creates high-quality vectorized images in PDF format for publication or distribution. Additionally, the browser accepts user-supplied input files such as BED files or GenomicRanges to display features or genomic interaction files that define co-accessibility, peak-to-gene linkages, or loops from chromatin conformation data.

To facilitate this interactive browser, ArchR utilizes the same optimizations described above for creating a genome-wide TileMatrix to create a TileMatrix for the chosen resolution specified within the plotting window. Cells corresponding to the same group are summed per tile and the resulting group matrix represents the accessibility in tiles across the specified window. This matrix can then be normalized by either the total number of reads in TSS/peak regions, the total number of cells, or the total number of unique nuclear fragments. By default, ArchR uses the reads in TSS regions, because this value is computed upon the creation of an Arrow file and is stable across analyses, unlike the peak regions. Because fragments in Arrow files are split per chromosome, the low memory cost and high speed of this process enables interactive visualization of hundreds of thousands of cells in seconds. Additionally, ArchR can plot tracks without the genome browser using the ArchRBrowserTrack function. ArchR also enables direct export of group normalized bigwig files using “export.bw” from Rtracklayer that can be directly used in conventional genome browsers.

### ArchR Methods - Peak Matrix

Once a peak set has been created (see ArchR Methods – Peak Calling), a cell x peak matrix can readily be made with ArchR. For each Arrow file, ArchR reads in scATAC-seq fragments from each chromosome and then computes overlaps with the peaks from the same chromosome. A sparse matrix cell x peak matrix is created for these peaks. The matrix is then added to the corresponding Arrow file. This procedure is iterated across each chromosome.

### ArchR Methods – Creation of Low-Overlapping Aggregates of Cells for Linkage Analysis

ArchR facilitates many integrative analyses that involve correlation of features. Performing these calculations with sparse single-cell data can lead to substantial noise in these correlative analyses. To circumvent this challenge, we adopted an approach introduced by Cicero^18^ to create low-overlapping aggregates of single cells prior to these analyses. We filter aggregates with greater than 80% overlap with any other aggregate in order to reduce bias. To improve the speed of this approach, we developed an implementation of an optimized iterative overlap checking routine and a implementation of fast feature correlations in C++ using the “Rcpp” package. These optimized methods are used in ArchR for calculating peak co-accessibility, peak-to-gene linkage, and for other linkage analyses.

### ArchR Methods – Peak Co-Accessibility

Co-accessibility analyses have been shown to be useful in downstream applications such as identifying groups of peaks that are all correlated forming “co-accessible networks”^18^. ArchR can rapidly compute peak co-accessibility from a peak matrix. These co-accessibility links can optionally be visualized using the ArchRBrowser. First, ArchR identifies 500+ low-overlapping cell aggregates (see Creation of Low-Overlapping Aggregates of Cells for Linkage Analysis). Second, for each chromosome (independently stored within an Arrow file), ArchR reads in the peak matrix and then creates the cell aggregate x peak matrix. ArchR next identifies all possible peak-to-peak combinations within a given window (by default 250 kb) and then computes the Pearson correlation of the log2-normalized cell aggregate x peak matrix. In this procedure, column sums across all chromosomes are used for depth normalization. ArchR iterates through all chromosomes and then combines the genome-wide results and stores them within the ArchRProject. These can be readily accessed for downstream applications. Additionally, ArchR enables users to lower the resolution of these interactions to better visualize the main interactors (keeping the highest correlation value observed in each window).

### ArchR Methods – Motif Annotations

ArchR enables rapid, fine-grained motif analyses. To carry out these analyses, ArchR must first identify the locations of all motifs in peak regions. ArchR natively supports access to motif sets curated from chromVAR^27^ and JASPAR^34^ to be used for these motif analyses. Additionally, ArchR makes possible the usage of multiple motif databases independently. ArchR first identifies motifs in peak regions using the matchMotifs function from the “motifmatchr” package (https://greenleaflab.github.io/motifmatchr/) with output being the motif positions within peaks. ArchR then creates a boolean motif overlap sparse matrix for each motif-peak combination that can be used for downstream applications such as enrichment testing and chromVAR. The motif positions and motif overlap matrix are stored on disk as an RDS file for later access, which minimizes the total memory of the ArchRProject, freeing memory for other analyses.

### ArchR Methods – Feature Annotations

ArchR allows for peak overlap analyses with defined feature sets. These feature sets could be ENCODE ChIP-seq/ATAC-seq peak sets or anything that can be specified as a GenomicRanges object. To facilitate this operation, we have curated a compendium of previously published ATAC-seq peak sets^19, 21–23, 26^, ENCODE ChIP-seq peak sets, and other custom feature sets for end-users^35^. We believe these custom feature sets will help users better annotate and describe cell types identified with scATAC-seq. These feature sets are overlapped with the ArchRProject peak set and then stored as a boolean feature overlap sparse matrix for each feature-peak combination that can be used for downstream applications such as enrichment testing and chromVAR. This feature overlap matrix is then stored on disk as an RDS file for later access, which minimizes the total memory of the ArchRProject, freeing memory for other analyses.

### ArchR Methods – Marker Peak Identification with Annotation Enrichment

ArchR allows for robust identification of features that are highly specific to a given group/cluster to elucidate cluster-specific biology. ArchR can identify these features for any of the matrices that are created with ArchR (stored in the Arrow files). ArchR identifies marker features while accounting for user-defined known biases that might confound the analysis (defaults are the TSS enrichment score and the number of unique nuclear fragments). For each group/cluster, ArchR identifies a set of background cells that match for the user-defined known biases and weights each equivalently using quantile normalization. Additionally, when selecting these bias-matched cells ArchR will match the distribution of the other user-defined groups. For example, if there were 4 equally represented clusters, ArchR will match the biases for a cluster to the remaining 3 clusters while selecting cells from the remaining 3 groups equally. By selecting a group of bias-matched cells, ArchR can directly minimize these confounding variables during differential testing rather than using modeling-based approaches. ArchR allows for binomial testing, Wilcoxon testing (via presto, https://github.com/immunogenomics/presto/), and two-sided t-testing for comparing the group to the bias-matched cells. These p-values are then adjusted for multiple hypothesis testing and organized across all group/clusters. This table of differential results can then be used to identify marker features based on user-defined log2(Fold Change) and FDR cutoffs.

### ArchR Methods – chromVAR Deviations Matrix

ArchR facilitates chromVAR analysis to identify deviation of accessibility within peak annotations (i.e. motif overlaps) compared to a controlled background set of bias-matched peaks. A challenge in using the published version of the chromVAR software is that it requires the full cell x peak matrix to be loaded into memory in order to compute these deviations. This can lead to dramatic increases in run time and memory usage for moderately sized datasets (∼50,000 cells). To circumvent these limitations, ArchR implements the same chromVAR analysis workflow by analyzing sample sub-matrices independently (see **Supplementary Fig. 12e**). First, ArchR reads in the global accessibility per peak across all cells. Second, for each peak, ArchR identifies a set of background peaks that are matched by GC-content and accessibility. Third, ArchR uses this background set of peaks and global accessibility to compute bias-corrected deviations with chromVAR for each sample independently. This implementation requires data from only 5,000-10,000 cells to be loaded into memory at any given time, minimizing the memory requirements, enabling scalable analysis with chromVAR, and improving run-time performance.

### ArchR Methods – Identification of Positive TF-Regulators

ATAC-seq allows for the unbiased identification of TFs that exhibit large changes in chromatin accessibility at sites containing their DNA binding motifs. However, families of TFs (for ex. GATA factors) share similar features in their binding motifs when looking in aggregate through position weight matrices (PWMs). This motif similarity makes it challenging to identify the specific TFs that might be driving observed changes in chromatin accessibility at their predicted binding sites. To circumvent this challenge, we have previously used gene expression to identify TFs whose gene expression is positively correlated to changes in the accessibility of their corresponding motif^21^. We term these TFs “positive regulators”. However, this analysis relies on matched gene expression data which may not be readily available in all experiments. To overcome this dependency, ArchR can identify TFs whose inferred gene scores are correlated to their chromVAR TF deviation scores. To achieve this, ArchR correlates chromVAR deviation scores of TF motifs with gene activity scores of TF genes from the low-overlapping cell aggregates (see above). When using scRNA-seq integration with ArchR, gene expression of the TF can be used instead of inferred gene activity score.

### ArchR Methods – TF Footprinting

ATAC-seq enables profiling of TF occupancy at base-pair resolution with TF footprinting. TF binding to DNA protects the protein-DNA binding site from transposition while the displacement or depletion of one or more adjacent nucleosomes creates increased DNA accessibility in the immediate flanking sequence. Collectively, these phenomena are referred to as the TF footprint. To accurately profile TF footprints, a large number of reads are required. Therefore, cells are grouped to create pseudo-bulk ATAC-seq profiles that can be then used for TF footprinting.

One major challenge with TF footprinting using ATAC-seq data is the insertion sequence bias of the Tn5 transposase^21, 36, 37^ which can lead to misclassification of TF footprints. To account for Tn5 insertion bias ArchR identifies the k-mer (user-defined length, default length 6) sequences surrounding each Tn5 insertion site. To do this analysis, ArchR identifies single-base resolution Tn5 insertion sites for each pseudo-bulk (see above Sample-Aware Pseudo-Bulk Replicate Generation), resizes these 1-bp sites to k-bp windows (-k/2 and + (k/2 - 1) bp from insertion), and then creates a k-mer frequency table using the “oligonucleotidefrequency(w=k, simplify.as=”collapse”)” function from the Biostrings package. ArchR then calculates the expected k-mers genome-wide using the same function with the BSgenome-associated genome file. To calculate the insertion bias for a pseudo-bulk footprint, ArchR creates a k-mer frequency matrix that is represented as all possible k-mers across a window +/- N bp (user-defined, default 250 bp) from the motif center. Then, iterating over each motif site, ArchR fills in the positioned k-mers into the k-mer frequency matrix. This is then calculated for each motif position genome-wide. Using the sample’s k-mer frequency table, ArchR can then compute the expected Tn5 insertions by multiplying the k-mer position frequency table by the observed/expected Tn5 k-mer frequency. For default TF footprinting with ArchR, motif positions (stored in the ArchRProject) are extended +/- 250 bp centered at the motif binding site. The pseudo-bulk replicates (stored as a HDF5-format coverage files) are then read into R as a coverage run-length encoding. For each individual motif, ArchR iterates over the chromosomes, computing a “Views” object using “Views(coverage,positions)”. ArchR uses an optimized C++ function to compute the sum per position in the Views object. This implementation enables fast and efficient footprinting **(Supplementary Fig. 12g-i).**

### ArchR Methods – Bulk ATAC-seq LSI projection

ArchR allows for projection of bulk ATAC-seq data into a scATAC-seq subspace as previously described^7^. ArchR first takes as input a bulk ATAC-seq sample x peak matrix and then identifies which peaks overlap the features used in the scATAC-seq dimensionality reduction. If there is sufficient overlap, ArchR estimates a scATAC-seq pseudo-cell x feature matrix within the features identified to overlap. These pseudo-cells (N = 250) per sample are sampled to be at 0.5x, 1x, 1.5x and 2x the average accessibility of the cell x feature matrix used. This step prevents unwanted sampling depth bias for this bulk projection analysis. The pseudo-cell x feature matrix is then normalized with the term-frequency x inverse document frequency (TF-IDF) method, using the same inverse document frequency obtained during the scATAC-seq dimensionality reduction. This normalized pseudo-cell x feature matrix is then projected with singular value decomposition “t(TF_IDF) %*% SVD$u %*% diag(1/SVD$d)” where TF_IDF is the transformed matrix and SVD is the previous SVD run using irlba in R. This reduced pseudo-cell x dim matrix can then be input to “uwot::umap_transform” which uses the previous scATAC-seq UMAP embedding to project the pseudo-cells into this embedding.

### ArchR Methods – Data Imputation with MAGIC

ArchR allows for using features such as gene scores and chromVAR deviation scores to assist in cluster annotation. However, features such as gene scores suffer from dropout noise in single-cell data. For scRNA-seq there have been many imputation methods developed to remedy this dropout noise. We have found that an effective method for imputation with scATAC-seq data is with Markov affinity-based graph imputation of cells (MAGIC)^38^. ArchR implements MAGIC for diffusing single-cell features across similar cells to smooth a single-cell matrix while simultaneously accounting for drop-out biases. MAGIC creates and stores a cell x cell diffusion matrix of weights that is then used to smooth the feature matrix with matrix multiplication. However, this diffusion matrix is dense and scales quadratically with the number of cells. To circumvent this limitation, ArchR creates equally sized blocks of cells (user-defined, default is 10,000) and then computes the partial diffusion matrix for these cells. These partial diffusion matrices are then combined to create a blocked diffusion matrix. This blocked diffusion matrix scales linearly in size leading to more memory efficiency but leads to lower resolution diffusion of data. To increase the resolution of this blocked diffusion matrix ArchR creates multiple replicates of the diffusion matrix to independently smooth the data matrix and then takes the average of the resulting smoothed matrices. ArchR additionally stores these blocked diffusion matrix replicates on-disk in HDF5-formatted files where each block is stored as its own group for direct access to specific parts of the matrix. ArchR’s MAGIC implementation shifts the memory usage to on-disk storage and thus enables data diffusion of extremely large datasets (N > 200,000) with minimal computing requirements.

### ArchR Methods – scATAC and scRNA Alignment

ArchR allows for efficient integration with scRNA-seq data utilizing Seurat’s integration infrastructure^11^. When performing this cross-platform alignment across large numbers of cells, we have found that the required memory and run time increase substantially. Moreover, constraining this alignment into smaller biologically relevant parts minimizes the alignment space into smaller alignment “sub-spaces”^7^. Thus, to increase alignment accuracy and improve runtime performance, ArchR enables the alignment of scATAC-seq and scRNA-seq to be constrained by user-defined groups of cells from both datasets that define smaller alignment sub-spaces. Within these sub-spaces, ArchR splits the scATAC-seq cells into equivalent slices of N cells (user-defined, default is 10,000 cells) and performs alignment with the scRNA-seq cells. This alignment procedure begins with the identification of the top variable genes (user-defined, default is 2,000 genes defined from scRNA-seq) using “Seurat::FindVariableFeatures”. Next, ArchR reads in the cell x gene scores matrix from the Arrow file for these cells. Then, ArchR imputes these gene scores using MAGIC and stores this imputed gene score matrix into a Seurat object for integration. ArchR then uses “Seurat::FindTransferAnchors” with canonical correlation analysis (CCA) to align this sub-space of cells efficiently. Next, ArchR extracts the aligned scRNA-seq cell, group, and gene expression profile with “Seurat::TransferData”. These gene expression profiles are stored in the corresponding Arrow files (stored as “GeneIntegrationMatrix”) for downstream analyses.

### ArchR Methods – scRNA Peak-To-Gene Linkage

We have previously used ATAC-seq peak-to-gene linkages to link putative enhancers and GWAS risk loci to their predicted target genes^7, 21^. ArchR can rapidly compute peak-to-gene links from a peak matrix and gene expression matrix (see above). These peak-to-gene links can optionally be visualized using the ArchRBrowser. First, ArchR identifies 500+ low-overlapping cell aggregates (see Creation of Low-Overlapping Aggregates of Cells for Linkage Analysis). Second, ArchR reads in the peak matrix and then creates the cell aggregate x peak matrix. Third, ArchR reads in the gene expression matrix and then creates the cell aggregate x gene matrix. ArchR then identifies all possible peak-to-gene combinations within a given window of the gene start (user-defined, default is 250 kb) and then computes the Pearson correlation of the log2-normalized cell aggregate x peak matrix and cell aggregate x gene matrix across all cell aggregates. ArchR computes these peak-to-gene links genome-wide and stores them within the ArchRProject, which can then be accessed for downstream applications. Additionally, ArchR enables users to lower the resolution of these interactions to better visualize the main interactors (keeping only the highest correlation value observed in each window).

### ArchR Methods – Cellular Trajectory Analysis

To order cells in pseudo-time, ArchR creates cellular trajectories that order cells across a lower N-dimensional subspace within an ArchRProject. Previously, we have performed this ordering in the 2-dimensional UMAP subspace^6^ but ArchR has improved upon this methodology to enable alignment within an N-dimensional subspace (i.e. LSI). First, ArchR requires a user-defined trajectory backbone that provides a rough ordering of cell groups/clusters. For example, given user-determined cluster identities, one might provide the cluster IDs for a stem cell cluster, then a progenitor cell cluster, and then a differentiated cell cluster that correspond to a known or presumed biologically relevant cellular trajectory (i.e. providing the cluster IDs for HSC, to MPP, to CMP, to Monocyte). Next, for each cluster, ArchR calculates the mean coordinates for each cell group/cluster in N-dimensions and retains cells whose Euclidean distance to those mean coordinates is in the top 5% of all cells. Next, ArchR computes the distance for each cell from clusteri to the mean coordinates of clusteri+1 along the trajectory and computes a pseudo-time vector based on these distances for each iteration of i. This allows ArchR to determine an N-dimensional coordinate and a pseudo-time value for each of the cells retained as part of the trajectory based on the Euclidean distance to the cell group/cluster mean coordinates. Next, ArchR fits a continuous trajectory to each N-dimensional coordinate based on the pseudo-time value using the “smooth.spline” function with df = 250 (degrees of freedom) and spar = 1 (smoothing parameter). Then, ArchR aligns all cells to the trajectory based on their Euclidean distance to the nearest point along the manifold. ArchR then scales this alignment to 100 and stores this pseudo-time in the ArchRProject for downstream analyses.

ArchR can create matrices that convey pseudo-time trends across features stored within the Arrow files. For example, ArchR can analyze changes in TF deviations, gene scores, or integrated gene expression across pseudo-time to identify regulators or regulatory elements that are dynamic throughout the cellular trajectory. First, ArchR groups cells in small user-defined quantile increments (default = 1/100) across the cellular trajectory. ArchR then smooths this matrix per feature using a user-defined smoothing window (default = 9/100) using the “data.table::frollmean” function. ArchR then returns this smoothed pseudo-time x feature matrix as a SummarizedExperiment for downstream analyses. ArchR additionally can correlate two of these smoothed pseudo-time x feature matrices using name matching (i.e. positive regulators with chromVAR TF deviations and gene score/integration profiles) or by genomic position overlap methods (i.e. peak-to-gene linkages) using low-overlapping cellular aggregates as described in previous sections. Thus, ArchR facilitates integrative analyses across cellular trajectories, revealing correlated regulatory dynamics across multi-modal data.

### Benchmarking Analysis – Preface

For benchmarking analyses, we used one of two computational environments: (1) a MacBook Pro laptop containing 32 GB of RAM and a 2.3GHz 8-core Intel Core i9 processor (16 threads) with data stored on an external USB hard drive; (2) a large-memory node on a high-performance cluster with 128 GB of RAM and two 2.40 GHz 10-core Intel Xeon E5-2640 V4 processors (20 threads). For benchmarking analyses using more limited compute resources (32 GB and 8 cores) we used the same large-memory node configuration but limited the available cores and memories using Slurm job submission properties. The main difference between the computational environment of the MacBook Pro and the server is the ability of each core on the MacBook Pro to use 2 threads whereas hyper-threading is disabled on the server and each core is effectively a single thread.

We downloaded scATAC-seq data from previously published and publicly available locations. We downloaded the immune cell data fragment files from Satpathy et al. 2019 (GSE129785), Granja et al. 2019 (GSE139369), and from the 10x Genomics website (https://www.10xgenomics.com/solutions/single-cell-atac/). For the mouse sci-ATAC-seq data, we downloaded the BAM files from http://atlas.gs.washington.edu/mouse-atac/. No additional steps were used prior to benchmarking analysis. We chose to focus our benchmarking tests versus Signac and SnapATAC based on the performance of LSI and LDM shown previously^9^. We ran all analyses in triplicate using snakemake via a slurm job submission engine on a high-performance cluster to accurately limit the available memory and cores. In the case of job failure, we allowed for multiple job attempts to ensure that analyses were reproducible. After each failed job attempt, the number of parallel threads for each software was lowered to attempt to complete the analysis without exceeding the available memory. Unless otherwise stated all analyses were run with default parameters for scATAC-seq benchmarking. We provide R markdown html files on our publication page https://github.com/GreenleafLab/ArchR_2020 detailing the exact procedures used for all benchmarking analyses.

### Benchmarking Analysis – Signac

Signac (https://github.com/timoast/signac) requires a predetermined peak set, thus we downloaded the previously published bulk hematopoiesis peak set from Corces et al. (ftp://ftp.ncbi.nlm.nih.gov/geo/series/GSE74nnn/GSE74912/suppl/GSE74912_ATACseq_All_Co unts.txt.gz) for all analyses. We first determined which cellular barcodes had more than 1,000 fragments by using “data.table::fread”. For each individual sample, we created a cell x peak matrix with the “FeatureMatrix” function using the fragment files and abundant cell barcodes as input. Then, we created a Seurat object from this cell x peak matrix with “CreateSeuratObject”. We then determined TSS enrichment scores for each cell across the first 3 chromosomes with the “TSSEnrichment” function. Default behavior for the TSSEnrichment function uses the first 2,000 TSSs; however, we increased this number (to include all TSSs on chr1-3) in order to stabilize the TSS enrichment scores for more consistent high-quality cell determination while still minimizing the run time. We then kept cells with a TSS enrichment score greater than 2 as high-quality cells passing filter. This TSS score cutoff differs from that of ArchR due to differences in the formula used for calculating TSS enrichment scores and differences in the gene annotation reference used by Signac. We then merged these individual Seurat objects (corresponding to each sample) and then performed TF-IDF normalization with “RunTFIDF” and “RunSVD” for LSI dimensionality reduction. We used the top 25% of features (ranked by accessibility) for LSI to reduce memory usage. The first 30 components were used by default for downstream analyses. Clusters were identified using “FindClusters” with default parameters. The scATAC-seq embeddings were determined using “RunUMAP” for UMAP and “RunTSNE” for tSNE respectively. Lastly, the gene score matrix was created using “FeatureMatrix” on the gene start and end coordinates (provided from ArchR) extended upstream by 2 kb for each sample and combined afterwards followed by log-normalization.

### Benchmarking Analysis - SnapATAC

SnapATAC (https://github.com/r3fang/SnapATAC) requires additional preprocessing steps prior to creation of a Snap file that can be used for downstream analyses. First, fragment files were sorted by their cell barcode with Unix “sort”. Next, these sorted fragment files were converted to Snap files by using SnapTools “snap-pre” with parameters “--min-mapq=30 --min-flen=50 --max-flen=1000 --keep-chrm=FALSE --keep-single=FALSE --keep-secondary=FALSE -- overwrite=TRUE --min-cov=1000 --max-num=20000 --verbose=TRUE” as described on the GitHub page. A genome-wide tile/bin matrix was then added using “snap-add-bmat” with parameters “--bin-size-list 5000” for a 5-kb matrix. To identify high-quality cells, SnapATAC computes a promoter ratio score for the fraction of accessible fragments that overlap promoter regions. We read in the 5-kb bin matrix into a Snap object using “addBmatToSnap” and then created a promoter Genomic Ranges object from the provided transcript annotation file (http://renlab.sdsc.edu/r3fang/share/github/reference/hg19/gencode.v30.annotation.gtf.gz) and then extending the gene start upstream by 2 kb. Next, we overlapped these regions using “findOverlaps” and then computed the summed accessibility within these overlapping regions vs the total accessibility across all 5-kb bins. We chose a cutoff for promoter ratio as 0.175 by manually inspecting the benchmarking dataset total accessibility vs promoter ratio plot as described in the GitHub. These high-quality cells were kept for downstream analyses. For dimensionality reduction, we first filtered bins that were greater than the 95^th^ percentile of non-zero bins. Next, we ran “runDiffusionMaps” with 30 eigenvectors to be computed (similar in the benchmarking analysis of all 3 methods). Clustering was performed with “runKNN” with the first 20 eigenvectors for a k-nn nearest neighbor search followed by “runCluster” with louvain.lib = “R-igraph”. The scATAC-seq embeddings were determined using “runViz” with method = “umap” for UMAP and method = “Rtsne” for tSNE for the top 20 eigenvectors. Lastly, the gene score matrix was determined by using the gene start and end coordinates (provided from ArchR) as input to “createGmatFromMat” with the input.mat = “bmat” and scaled with “scaleCountMatrix”. For comparing estimated dimensionality reduction in SnapATAC (estimated LDM) to estimated LSI in ArchR, we first sampled N cells (10,000 or the number of cells specified) based on the inverse of their coverage and then computed diffusion maps with “runDiffusionMaps”. The remaining cells were projected with “runDiffusionMapsExtension” and the two Snap objects were combined for downstream analysis.

### Benchmarking Analysis - ArchR

For analysis with ArchR, we first converted input scATAC-seq data (fragment files or BAM files) to Arrow files with “createArrowFiles” with minFrags = 1000, filterTSS = 4, and addGeneScoreMat = FALSE (addGeneScoreMat was set to false to allow for downstream benchmarking of this individual step). These Arrow files were then used to create an ArchRProject with the appropriate genome annotation. We identified doublet scores for each sample with “addDoubletScores” and “filterDoublets” respectively; however, time and memory used for doublet identification were not included in the benchmarking results because this step is unique to ArchR and would complicate direct comparisons to other software. We then computed the iterative LSI dimensionality reduction with “addIterativeLSI” with default parameters (variableFeatures = 25,000 and iterations = 2). Clusters were identified using “addClusters” with default parameters. The scATAC-seq embeddings were determined using “addUMAP” for UMAP and “addTSNE” for tSNE. Lastly, the gene score matrix was added by “addGeneScoreMatrix” which stores the depth-normalized cell x gene matrix. For comparison of estimated LSI in ArchR to estimated LDM in SnapATAC, “addIterativeLSI” was run with an additional parameter for sampling (sampleCellsFinal = 10,000 or the number of cells specified).

### ArchR Analysis – Comparison of Gene Score Methods

We used ArchR to benchmark 53 models of inferring gene scores to emulate gene expression. All models were tested with the same gene annotation reference for direct comparison. We additionally used Signac, SnapATAC, and co-accessibility to create gene score models for comparison, making a total of 56 models. We used two datasets for evaluation: (1) ∼30,000 PBMCs and (2) ∼30,000 bone marrow cells. We first created the gene score models that incorporated distance by systematically changing the input parameters for “addGeneScoreMatrix”. This parameter sweep included TSS exponential decay functions (useTSS = TRUE) and gene body exponential decay functions (useTSS = FALSE). We tried other decay functions but saw no appreciable difference so we used exponential decay (this is a user-input so any model as a function of relative distance may be inserted). For gene score models that were overlap-based (no distance function), we used “addFeatureMatrix” based on a set of genomic regions corresponding to either an extended gene promoter [resize(genes, 1, “start”) followed by resize(2*window + 1, “center”)] or an extended gene body [extendGR(genes, upstream, downstream)]. For each model, we created a genome-wide gene score matrix and extracted these matrices from the Arrow files using “getMatrixFromProject”. We next created 500 low-overlapping random groupings of 100 cells with ArchR (see above) and took the average gene scores for each of these groupings. Next, we collected the gene scores calculated by Signac and SnapATAC during our benchmarking tests and averaged the gene scores across the same groupings. For co-accessibility, we created gene scores as previously described with Cicero^6, 7, 18^. We first used Cicero to create 5,000 lowly-overlapping cell groupings of 50 cells with “cicero_cds”. Next, we calculated the average accessibility for these groupings across all peaks (with getMatrixFromProject). We correlated all peaks within 250 kb to get peak co-accessibility. We annotated the peaks as promoter if within 2.5 kb from the gene start with “annotate_cds_by_site”. Finally, gene scores for the co-accessibility model were identified with “build_gene_activity_matrix” with a co-accessibility cutoff of 0.35 followed by “normalize_gene_activities”. For this co-accessibility model, we tested various parameters such as promoter window size, correlation cutoff, and peak-to-peak distance maximums to make sure the results were reproducible.

Having a cell aggregate x gene score matrix for all 56 models, we next created a gene expression matrix to test these models. We integrated our scATAC-seq (from ArchR’s results) with previously annotated scRNA-seq datasets (10k PBMC from 10x website and Bone Marrow from Granja et al., 2019) using “Seurat::FindTransferAnchors” and “Seurat::TransferData” with the top 2,000 variable genes from scRNA-seq. This integration was performed for each scATAC-seq sample independently and the scRNA-seq data used for each bone marrow alignment was constrained to match cell sources together (i.e. BMMC scATAC-seq with BMMC scRNA-seq and CD34+ scATAC-seq with CD34+ scRNA-seq)^7^. From this integration, each scATAC-seq cell was paired to a matched gene expression profile. We averaged the gene expression profiles for each of the 500 lowly-overlapping groups (see above) to create a cell aggregate x gene expression matrix.

To benchmark the performance for each gene score model, we identified 2 gene sets: the top 2,000 variable genes defined by “Seurat::FindVariableGenes” and the top 1,000 differentially expressed genes defined by “Seurat::FindAllMarkers” (ranking the top N genes for each scRNA-seq cluster until 1,000 genes were identified). For these gene sets, we calculated the gene-wise correlation (how well do the gene score and gene expression correlate across all genes) and the aggregate-wise correlation (how well do the gene score and gene expression correlate across all cell aggregates). These 4 measures were then ranked across all models, and the average ranking was used to score the 56 models.

To orthogonally support this result, we downloaded previously published paired bulk ATAC-seq + RNA-seq for hematopoiesis^19^. We then iteratively down-sampled the reads from each dataset to create 100 pseudo-cells with 10,000 fragments from each bulk ATAC-seq sample. We then created a scATAC-seq fragments file for each pseudo-cell. We performed an identical analysis as described above for the 53 ArchR gene score models. For comparing these 53 models, we used 2 gene sets: the top 2,000 variable genes defined by log2-normalized expression-ranked variance across each cell type and the top 1,000 marker genes defined by the top log2(fold change) for each cell type vs the average expression of all cell types. We similarly ranked the gene-wise and aggregate-wise correlation across all models, and used the average ranking to score each model.

### ArchR Analysis – Large Simulated PBMC ∼1.2M Cells

To further test ArchR’s capability to analyze extremely large datasets (N > 200,000), we simulated ∼1.3M single cells contained within 200 fragment files. We used 4 PBMC samples (2 x 5,000 cells and 2 x 10,000 cells from 10x Genomics) for creating this large dataset. We randomly shifted each scATAC-seq fragment with a mean difference of +/- 50-100 bp (randomly sampled) and a standard deviation of +/- 10-20 bp (randomly sampled). We then sampled the fragments by 80% to ensure some differences between simulated cells and then saved these to bg-zipped fragment files. We then used ArchR to convert these fragment files to Arrow files with “createArrowFiles” with minFrags = 1000, filterTSS = 4 and addGeneScoreMat = TRUE. These Arrow files were then assembled into an ArchRProject. We identified doublet scores for each simulated dataset with “addDoubletScores” and “filterDoublets” respectively, retaining ∼1.2 million cells after doublet removal. We then computed the estimated iterative LSI dimensionality reduction with “addIterativeLSI” (variableFeatures = 25,000, sampleCellsFinal = 25,000 and 2 iterations). Estimated clusters were identified using “addClusters” with sampleCells = 50,000. This estimation method uses a subset of cells to cluster and then the remaining cells are annotated by their nearest neighbors (the maximum annotation observed). An estimated scATAC-seq UMAP was created using “addUMAP” with sampleCells = 100,000. This estimation method uses a subset of cells to create a UMAP embedding and then the remaining cells are projected into the single-cell embedding using “umap::umap_transform”.

### ArchR Analysis – Large Hematopoiesis 220K Cells

We wanted to test ArchR’s full analysis suite with a large dataset (N > 200,000) comprised of previously published immune cell data^6, 7^. We additionally grouped all Fluidigm C1-based scATAC-seq data from Buenrostro et al. 2018^4^ into a fragment file. This amounted to a total of 49 scATAC-seq fragment files corresponding to over 200,000 cells. We first used ArchR to convert these fragment files to Arrow files using “createArrowFiles” with minFrags = 1000, filterTSS = 8 and addGeneScoreMat = TRUE. These Arrow files are then used to create an ArchRProject. We identified doublet scores for each simulated dataset with “addDoubletScores” and “filterDoublets” respectively. We then computed the estimated iterative LSI dimensionality reduction with “addIterativeLSI” (variableFeatures = 25,000, sampleCellsFinal = 25,000 and iterations = 2). A scATAC-seq UMAP was then created by using “addUMAP” with minDist = 1 and nNeighbors = 40. Clusters were initially identified using “addClusters” with default parameters. We re-clustered the early progenitor cells (clusters containing CD34+ cells) with a clustering resolution of 0.4 to better resolve these cell clusters. We added MAGIC imputation weights with “addImputationWeights” for imputing single-cell features that are then overlaid on the UMAP embedding. We then manually merged and assigned clusters that correspond to cell types based on known marker gene scores and observation of sequencing tracks using the ArchRBrowser.

To identify a union peak set, we created group coverage files, which contain the aggregated accessibility of groups of single cells within a cluster, with “addGroupCoverages”. We then created a reproducible peak set with “addReproduciblePeakSet” and a cell x peak matrix with “addPeakMatrix”. Next, we determined background peaks that are matched in GC-content and accessibility with “addBgdPeak”. For downstream motif-based analyses we added motif overlap annotations with “addMotifAnnotations” for CIS-BP version 1 motifs (version = 1). We computed a ChromVAR deviations matrix with “addDeviationsMatrix”. We next identified positive TF regulators with “correlateMatrices” where useMatrix1 = “MotifMatrix” and useMatrix2 = “GeneScoreMatrix”. To identify which of these correlated TF regulators had strong differential motif activity differences we calculated the average motif deviation scores with “exportGroupSE” for each cluster and computed the max observed deviation difference between any two clusters. This motif difference and the TF-to-gene score correlation were then used to identify positive regulators (correlation > 0.5 and a maximum deviation score difference > 50^th^ percentile). Differential accessibility for each cluster was determined using “markerFeatures” with maxCells = 1000 and useMatrix = “PeakMatrix”. Marker peaks were defined as peaks with a log2(Fold Change) > 1.5 and an FDR < 0.01 (Wilcoxon-test with presto, https://github.com/immunogenomics/presto/). We then determined enriched motifs with “peakAnnoEnrichment” in these marker peaks and plotted the motif enrichment p-values for the positive TF regulators. ArchR has a curated set of previously published bulk ATAC-seq datasets that we used for feature overlap enrichment by computing overlaps with “addArchRAnnotations” (collection = “ATAC”) and “peakAnnoEnrichment”. TF footprints, with Tn5-bias correction, were calculated by “plotFootprints” with motif positions from “getPositions” and normMethod = subtract. Bulk hematopoietic ATAC-seq (GSE74912) was projected into the scATAC-seq subspace using “projectBulkATAC” with N = 250 cells. Peak co-accessibility was computed with “addCoAccessibility” and accessibility tracks were created with the ArchRBrowser.

We next wanted to integrate our scATAC-seq data with previously published hematopoietic scRNA-seq data^7^. To do this analysis, we used “addGeneIntegrationMatrix” with sampleCellsATAC = 10,000, sampleCellsRNA = 10,000, and a groupList specifying to group cells from T/NK clusters and cells from non-T/NK clusters for both scATAC-seq and scRNA-seq prior to alignment. This constrained integration improved the alignment accuracy and added a matched gene expression profile for each scATAC-seq cell. We overlaid these gene expression profiles on the UMAP embedding with “plotEmbedding”. After this integration analysis, we identified peak-to-gene links with “addPeak2GeneLinks” and visualized them with “peak2GeneHeatmap”.

To create a cellular trajectory across B cell differentiation, we used “addTrajectory” with preFilterQuantile = 0.8, useAll = FALSE, and an initial trajectory of “HSC -> CMP.LMPP -> CLP.1 -> CLP.2 -> PreB -> B”. We next created trajectory matrices for “MotifMatrix”, “GeneScoreMatrix”, “GeneIntegrationMatrix” and “PeakMatrix”. We correlated the deviation score and gene score trajectory matrices with “correlateTrajectories”. Additionally, we correlated the deviation score and gene expression trajectory matrices with “correlateTrajectories”. We kept TFs whose correlation was 0.5 or greater for both of the correlation analyses. We determined these TFs as positive TF regulators across the B cell trajectory. We also used ArchR to identify peak-to-gene links across the B cell trajectory with “correlateTrajectories” with useRanges = TRUE, varCutOff1 = 0.9, and varCutOff2 = 0.9. Lastly, we grouped cells into 5 groups of cells based on pseudo-time across the B cell trajectory for track visualization (with the ArchRBrowser) and TF footprinting of the TF regulators.

## ACKNOWLEDGEMENTS

We thank members of the Greenleaf and Chang laboratories for helpful comments. This work was supported by NIH RM1-HG007735 and UM1-HG009442 (to H.Y.C. and W.J.G.), R35-CA209919 (to H.Y.C.), UM1-HG009436 and U19-AI057266 (to W.J.G.), K99-AG059918 and the American Society of Hematology Scholar Award (to M.R.C.), and an International Collaborative Award (to H.Y.C., H.C.).

## AUTHOR CONTRIBUTIONS

J.M.G., M.R.C., H.Y.C. and W.J.G conceived the project. J.M.G. and M.R.C. led the design of the ArchR software with input from S.E.P and W.J.G. M.R.C. led the scATAC-seq data creation with input from S.T.B., H.C. and H.Y.C.. J.M.G. and M.R.C. led the single-cell analysis presented in this paper. J.M.G., M.R.C., H.Y.C. and W.J.G wrote the manuscript with input from all authors.

## Supplementary Figure Legends

**Supplementary Fig. 1.**
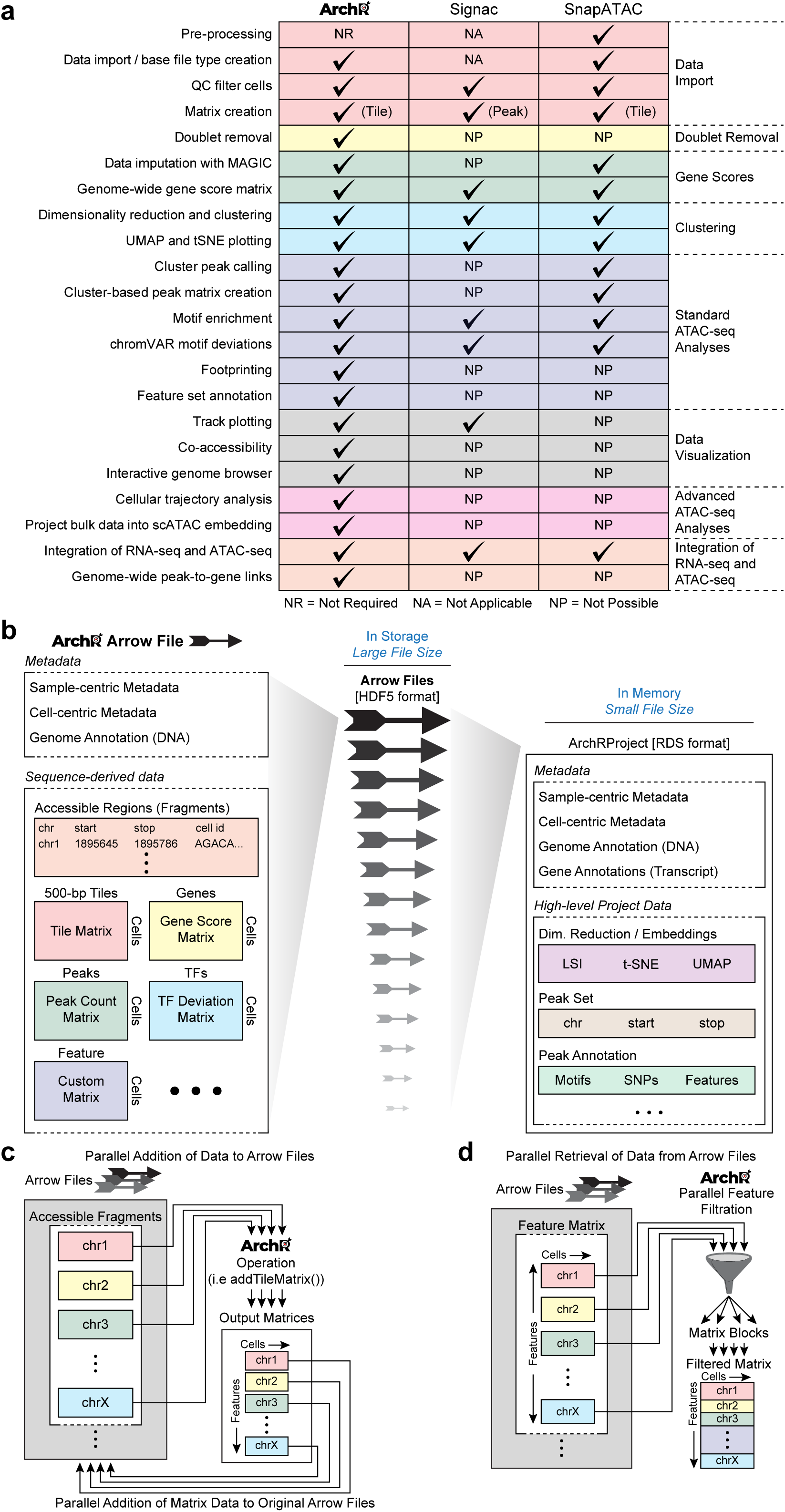
ArchR infrastructure and supported analyses. **a.** Comparison of supported scATAC-seq analysis features across ArchR, Signac and SnapATAC. **b.** (Left) Schematic of the ArchR Arrow file format where <u>a</u>ccessible reads and arrays are <u>o</u>rganized <u>w</u>ithin. Arrow files can then be used as input for an ArchRProject (Right). The ArchRProject stores the locations of these Arrow files and extracts their cell-centric metadata. All analysis with ArchR operates through this ArchRProject which can readily access data from Arrow files stored on disk. **c.** Schematic demonstrating how ArchR operations that involve using Arrow fragments (i.e. addTileMatrix) operate on each chromosome independently in parallel for many Arrow files and then add the resulting matrix back to the corresponding Arrow files again in parallel. **d.** Schematic demonstrating how ArchR operations that use Arrow matrices (i.e. addIterativeLSI) access a subset of each chromosome’s matrix from each Arrow file in parallel that are then merged to create a filtered matrix for subsequent analysis.

**Supplementary Fig. 2.**
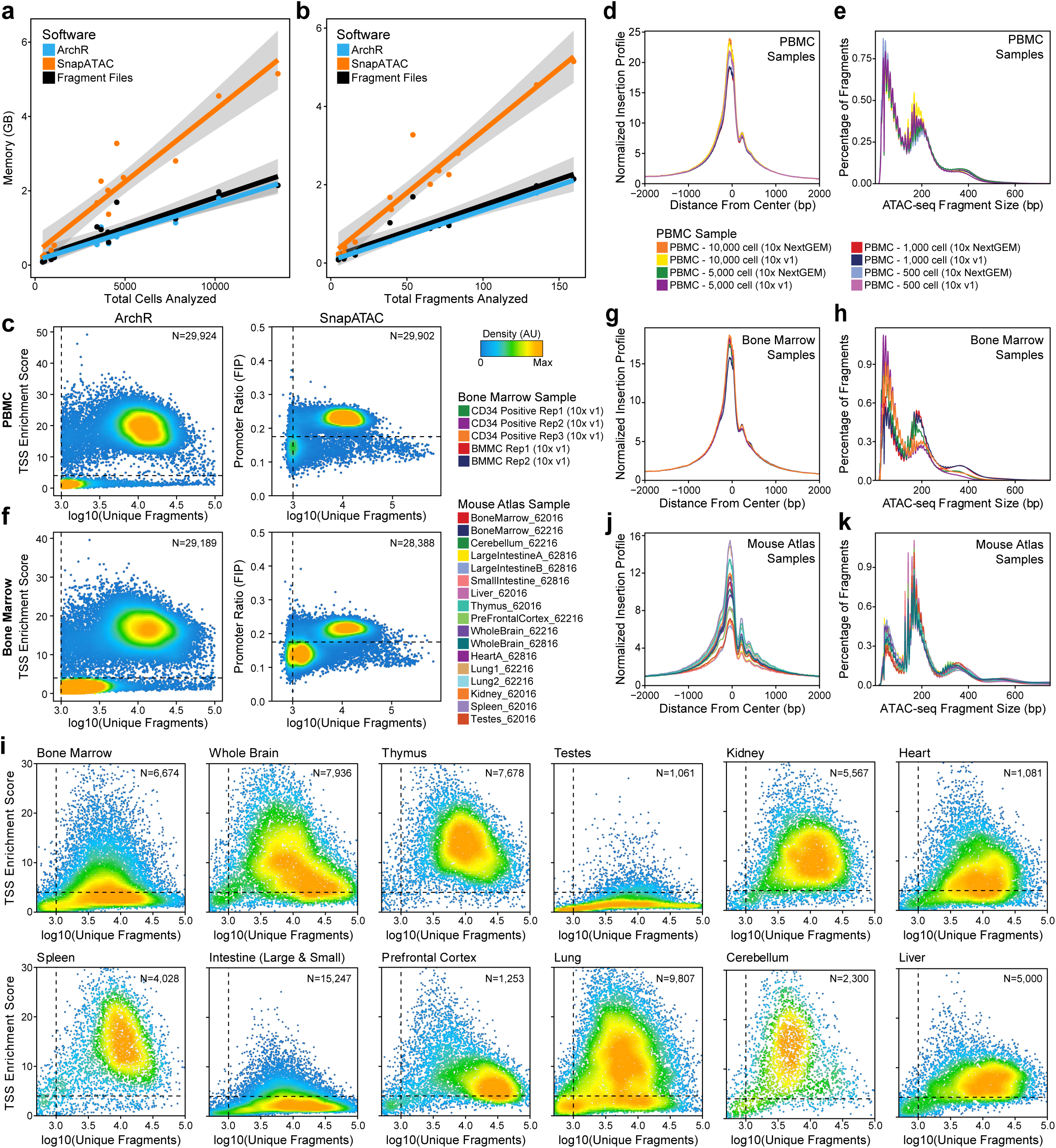
**a-b.** File sizes of storage formats (for both accessible fragments and counts matrix) for ArchR and SnapATAC compared to (**a**) the total number of cells they represent or (**b**) the total number of fragments corresponding to the cells represented in each file. Line colors represent the different software used or the original fragment files. **c.** QC filtering plots for the PBMCs dataset from (left) ArchR, showing the TSS enrichment score vs unique nuclear fragments per cell, or (right) SnapATAC, showing the promoter ratio / fraction of reads in promoters (FIP) vs unique nuclear fragments per cell. Dot color represents the density in arbitrary units of points in the plot. **d-e.** Aggregate (**d**) TSS insertion profiles centered at all TSS regions or (**e**) fragment size distributions for the cells passing ArchR QC thresholds for each sample in the PBMCs dataset. Line color represents the sample from the dataset as indicated below the plot. **f.** QC filtering plots for the bone marrow cell dataset from (left) ArchR, showing the TSS enrichment score vs unique nuclear fragments per cell, or (right) SnapATAC, showing the promoter ratio / fraction of reads in promoters (FIP) vs unique nuclear fragments per cell. Dot color represents the density in arbitrary units of points in the plot. **g-h.** Aggregate (**g**) TSS insertion profiles centered at all TSS regions or (h) fragment size distributions for the cells passing ArchR QC thresholds for each sample in the bone marrow cell dataset. Line color represents the sample from the dataset as indicated below the plot. **i.** QC filtering plots from ArchR for each individual organ type from the mouse atlas dataset showing the TSS enrichment score vs unique nuclear fragments per cell. Dot color represents the density in arbitrary units of points in the plot. **j-k.** Aggregate (**j**) TSS insertion profiles centered at all TSS regions or (**k**) fragment size distributions for the cells passing ArchR QC thresholds for each sample in the mouse atlas dataset. Line colors represent different samples as indicated to the left of the plot.

**Supplementary Fig. 3.**
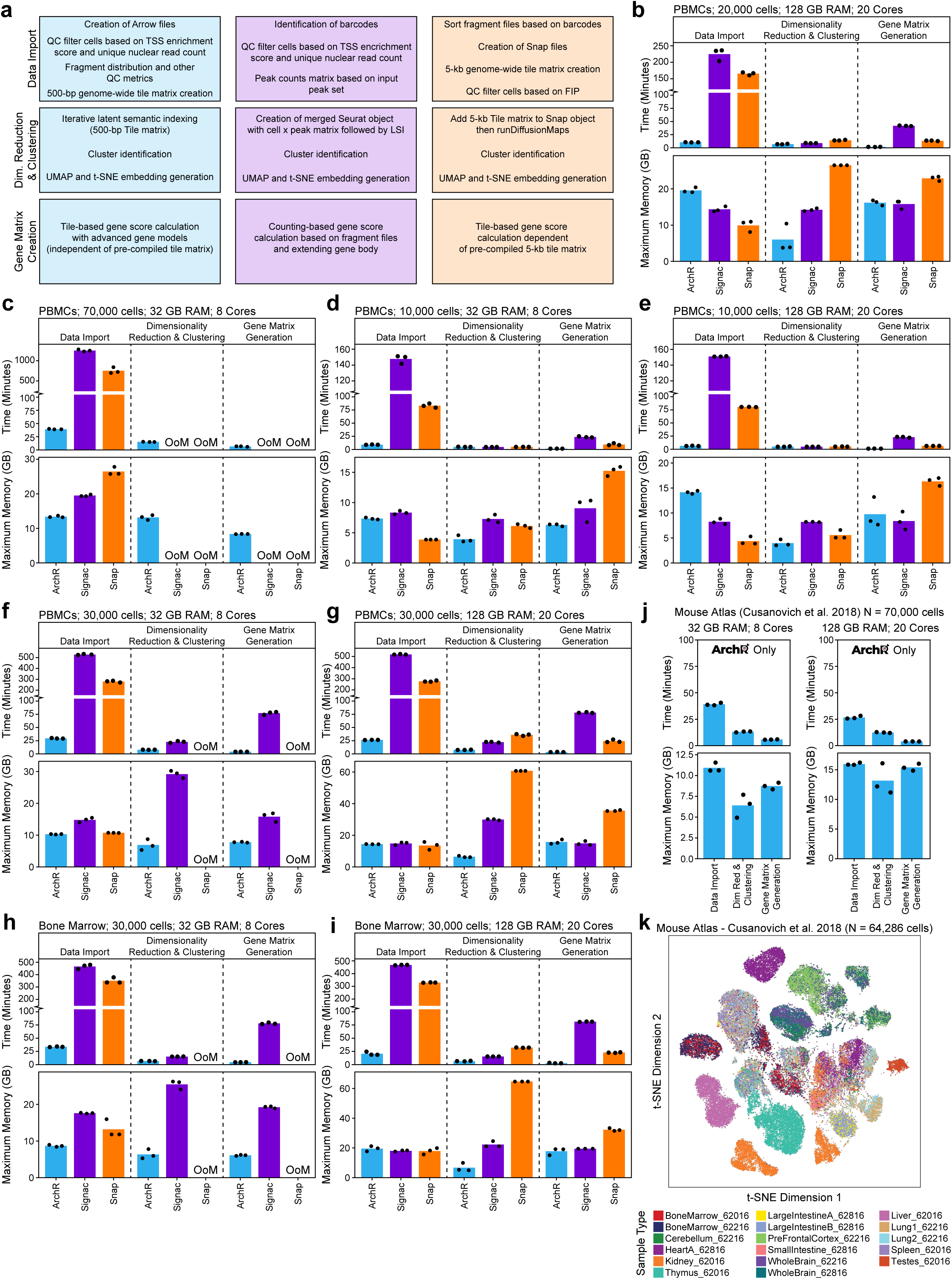
**a.** Schematic describing the individual benchmarking steps compared across ArchR, Signac, and SnapATAC for (1) Data Import, (2) Dimensionality Reduction and Clustering, and (3) Gene Score Matrix Creation. **b-i.** Comparison of ArchR, Signac, and SnapATAC for run time and peak memory usage for the analysis of (**b**) ∼20,000 cells from the PBMCs dataset using 128 GB of RAM and 20 cores (plot corresponds to Figure 1b), (**c**) ∼70,000 cells from the PBMCs dataset using 32 GB of RAM and 8 cores (plot corresponds to Figure 1c), (**d-e**) ∼10,000 cells from the PBMCs dataset using (**d**) 32 GB of RAM and 8 cores or (**e**) 128 GB of RAM and 20 cores, (**f-g**) ∼30,000 cells from the PBMCs dataset using (**f**) 32 GB of RAM and 8 cores or (**g**) 128 GB of RAM and 20 cores, and (**h-i**) ∼30,000 cells from the bone marrow dataset using (**h**) 32 GB of RAM and 8 cores or (**i**) 128 GB of RAM and 20 cores. Dots represent individual replicates of benchmarking analysis. **j.** Benchmarks from ArchR for run time and peak memory usage for the analysis of ∼70,000 cells from the sci-ATAC-seq mouse atlas dataset for (left) 32 GB of RAM with 8 cores and (right) 128 GB of RAM with 20 cores. Dots represent individual replicates of benchmarking analysis. **k.** t-SNE of mouse atlas scATAC-seq data (N = 64,286 cells) colored by individual samples.

**Supplementary Fig. 4.**
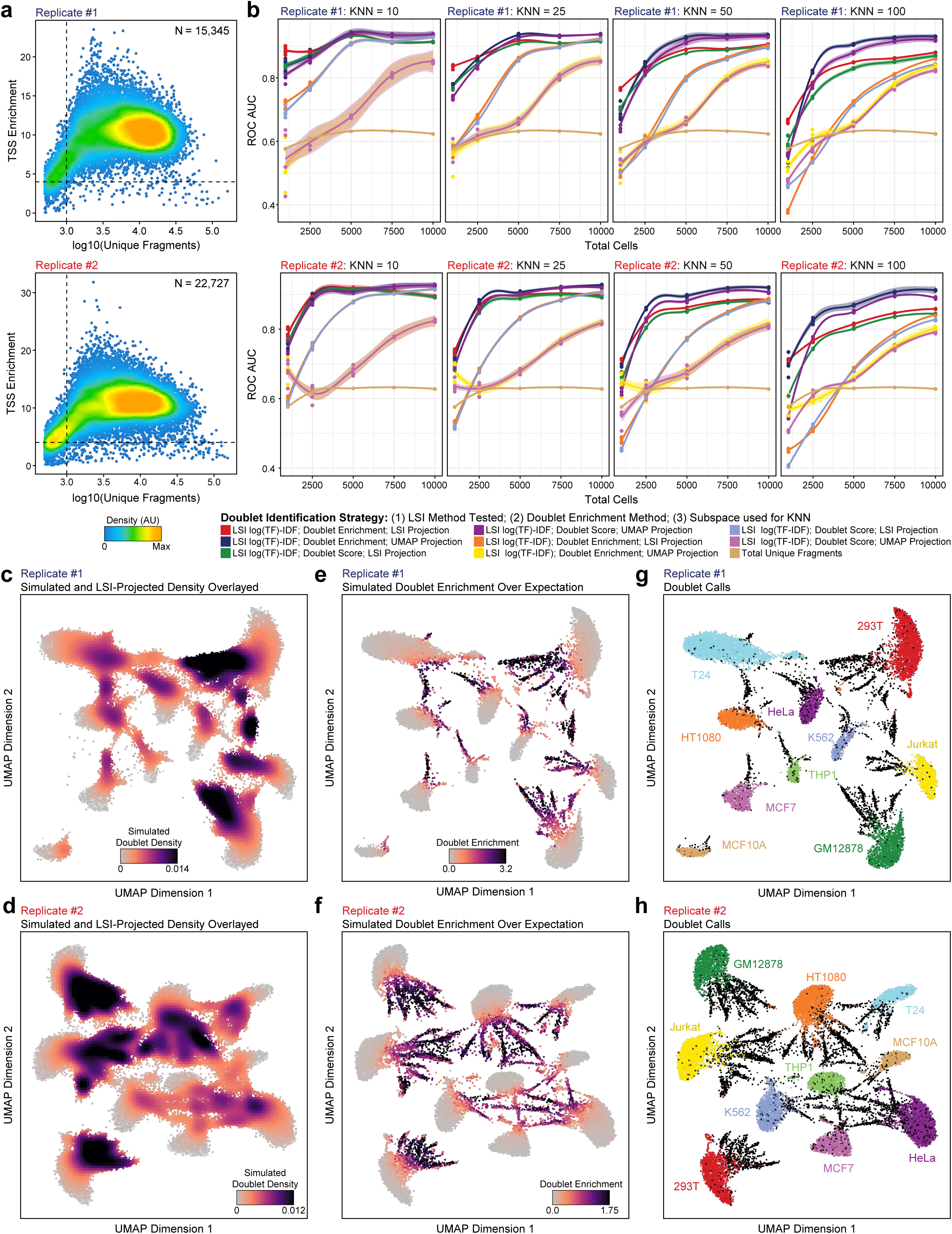
**a.** QC filtering plots from ArchR for (top) replicate 1 and (bottom) replicate 2 from the cell line mixing dataset showing the TSS enrichment score vs unique nuclear fragments per cell. Dot color represents the density in arbitrary units of points in the plot. **b.** Accuracy of various doublet prediction methods for (top) replicate 1 and (bottom) replicate 2 from the cell line mixing dataset, measured by the area under the curve (AUC) of the receiver operating characteristic (ROC), across different in silico cell loadings. Accuracy is determined with respect to genotype-based identification of doublets using demuxlet. Above each plot, “KNN” represents the number of cells nearby each projected synthetic doublet to record when calculating doublet enrichment scores. The distance for KNN recording is determined in the LSI subspace for LSI projection and in the UMAP embedding for UMAP projection parameters. **c-h.** UMAP of scATAC-seq data showing the (**c-d**) simulated doublet density, (**e-f**) simulated doublet enrichment, or (**g-h**) cell line identity based on genotyping information and demuxlet for (**c,e,g**) replicate 1 (N = 15,345 cells) and (**d,f,h**) replicate 2 (N = 22,727 cells) of the cell line mixing dataset.

**Supplementary Fig. 5.**
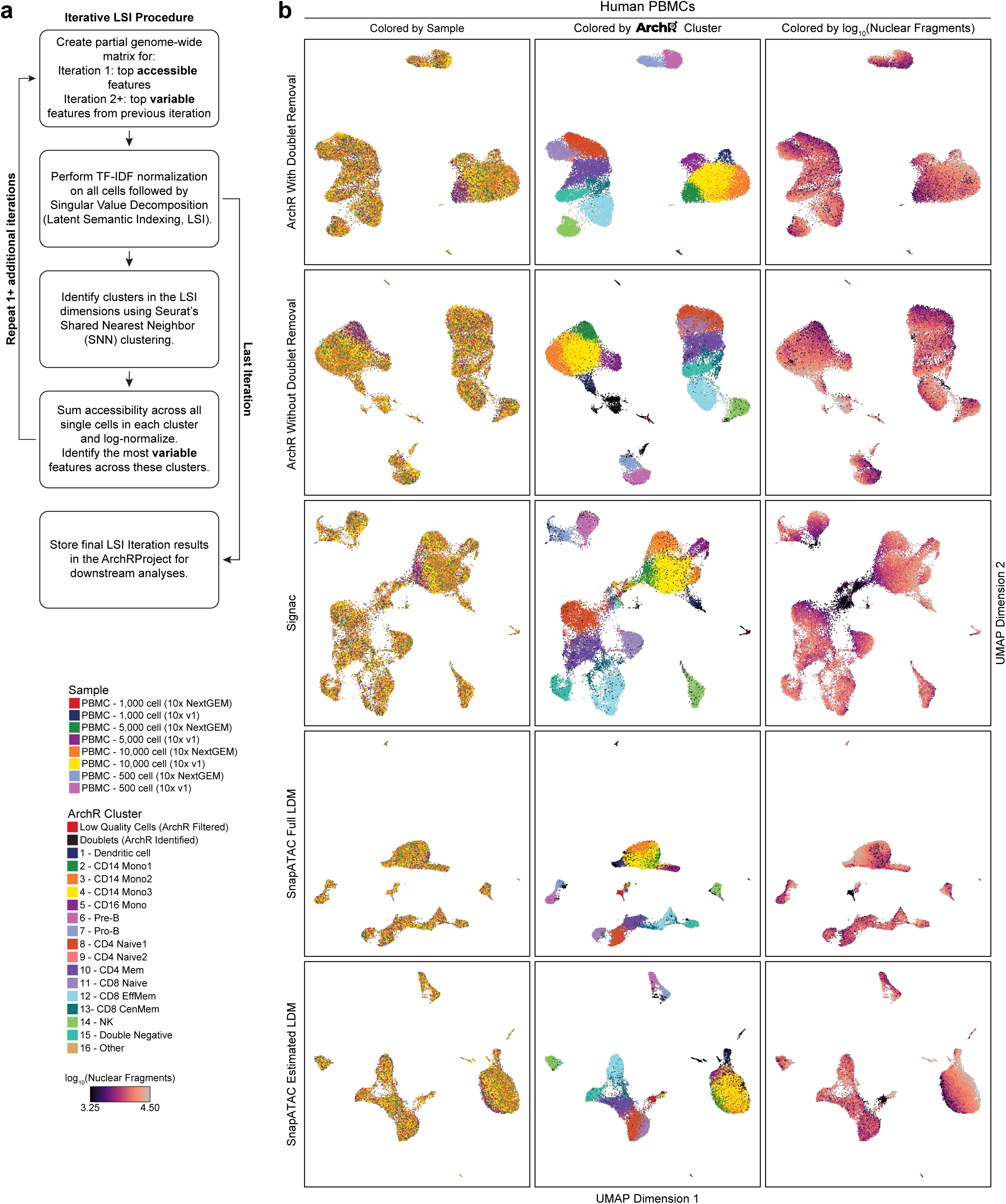
**a.** Schematic of the iterative LSI procedure implemented in ArchR for dimensionality reduction. **b.** UMAPs of scATAC-seq data from ∼30,000 cells from the PBMCs dataset to compare clustering results across ArchR with doublet removal, ArchR without doublet removal, Signac, SnapATAC, and SnapATAC with estimated LDM. Each UMAP is colored by (left) sample, (middle) clusters as defined by ArchR with doublet removal, and (right) the number of unique nuclear fragments.

**Supplementary Fig. 6.**
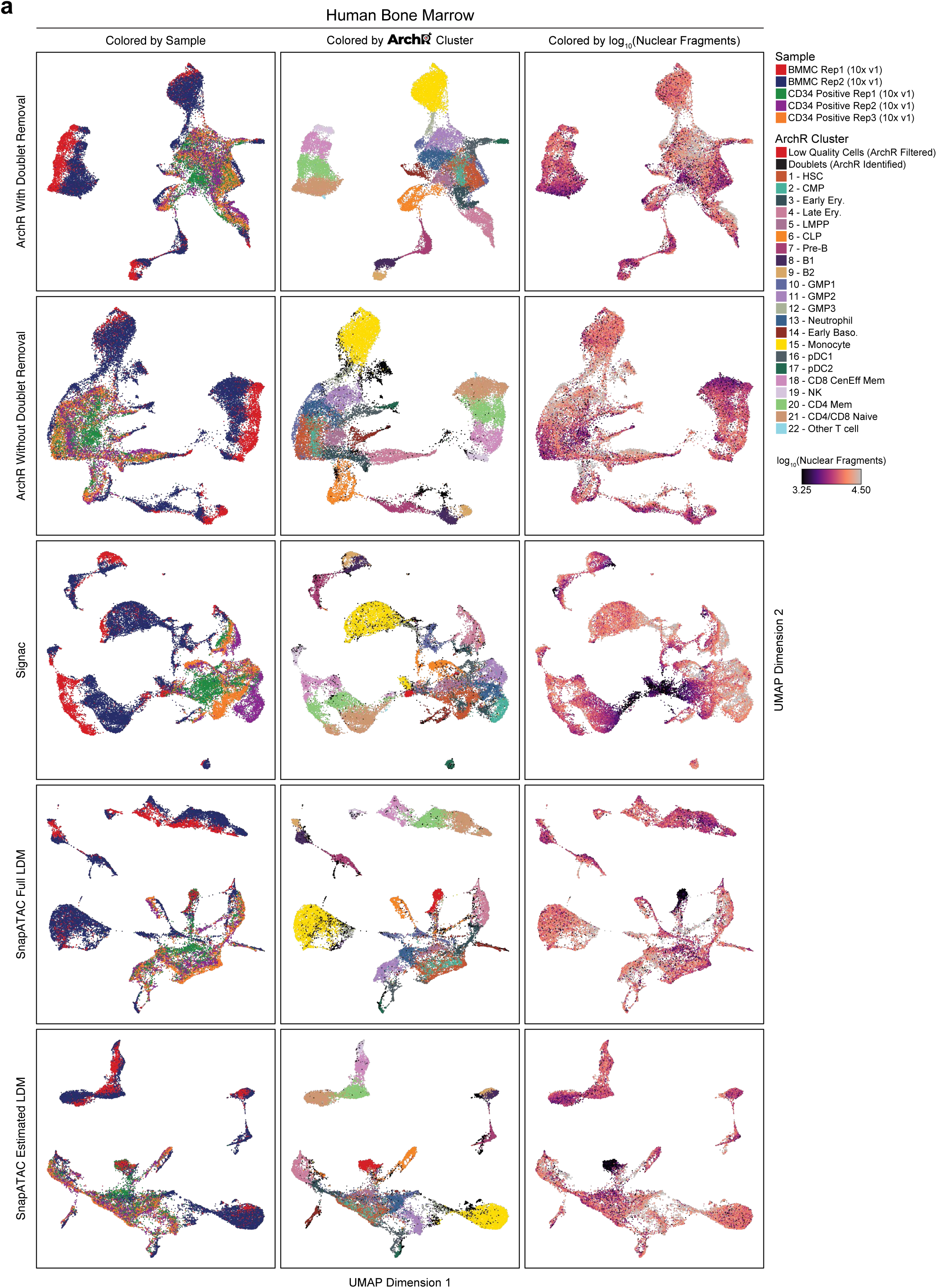
**a.** UMAPs of scATAC-seq data from ∼30,000 cells from the bone marrow dataset to compare clustering results across ArchR with doublet removal, ArchR without doublet removal, Signac, SnapATAC, and SnapATAC with estimated LDM. Each UMAP is colored by (left) sample, (middle) clusters as defined by ArchR with doublet removal, and (right) the number of unique nuclear fragments.

**Supplementary Fig. 7.**
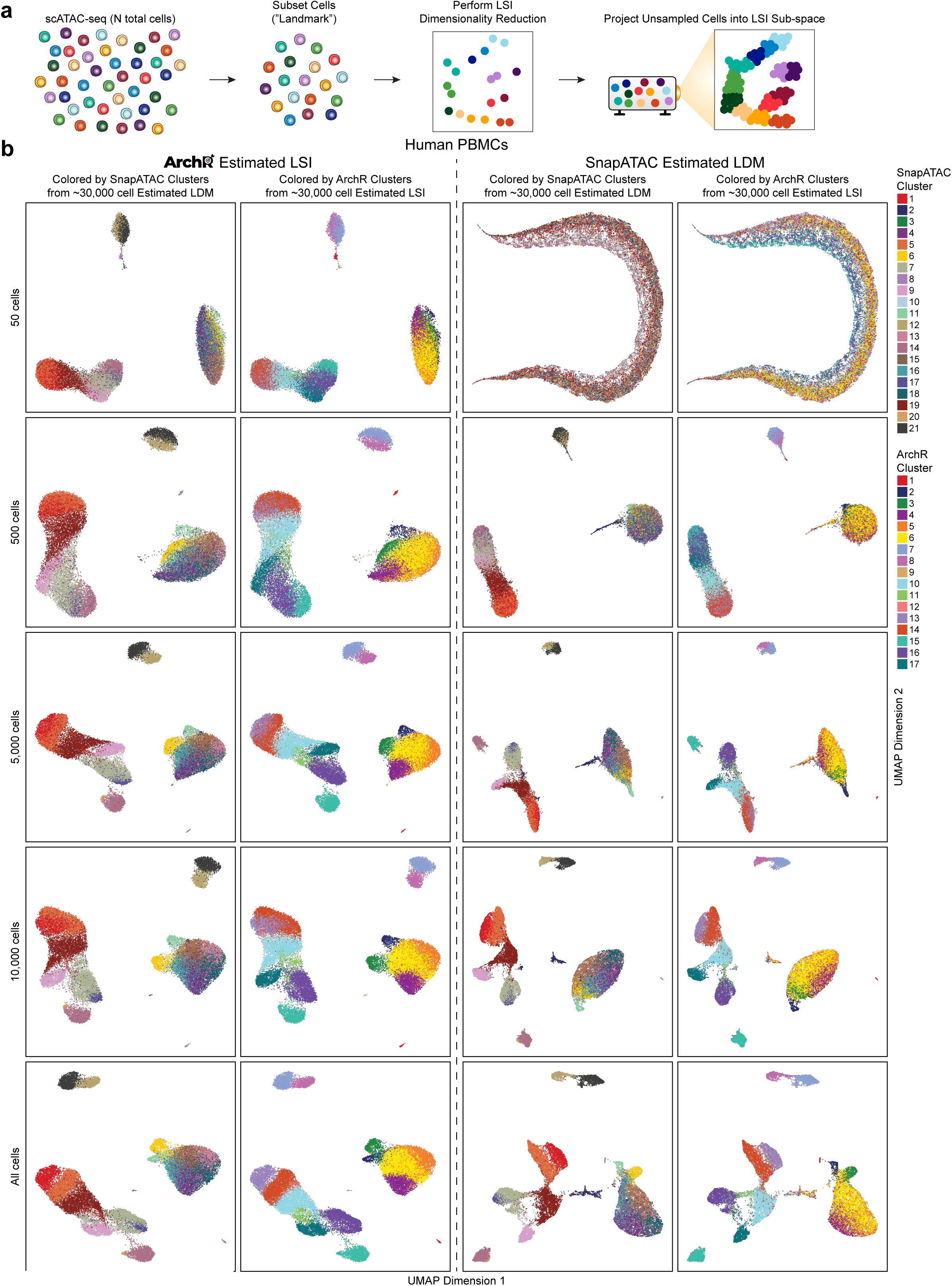
**a.** Schematic of the estimated LSI framework implemented by ArchR. Briefly, a subset of cells, referred to as “landmark” cells, are used for LSI dimensionality reduction. The remaining cells are then linearly projected with LSI projection into this landmark-defined LSI subspace. This method enables massive-scale analysis of scATAC-seq data with ArchR. **b.** UMAPs of scATAC-seq data from ∼30,000 cells from the PBMCs dataset showing the results of dimensionality reduction from (left) estimated LSI with ArchR after doublet removal or (right) estimated LDM with SnapATAC. For each analytical case, a range of cell numbers is used for the landmark cell subset (top to bottom). Within each analytical case, two UMAPs are presented, colored by the clusters identified without estimation from (left) ArchR or (right) SnapATAC.

**Supplementary Fig. 8.**
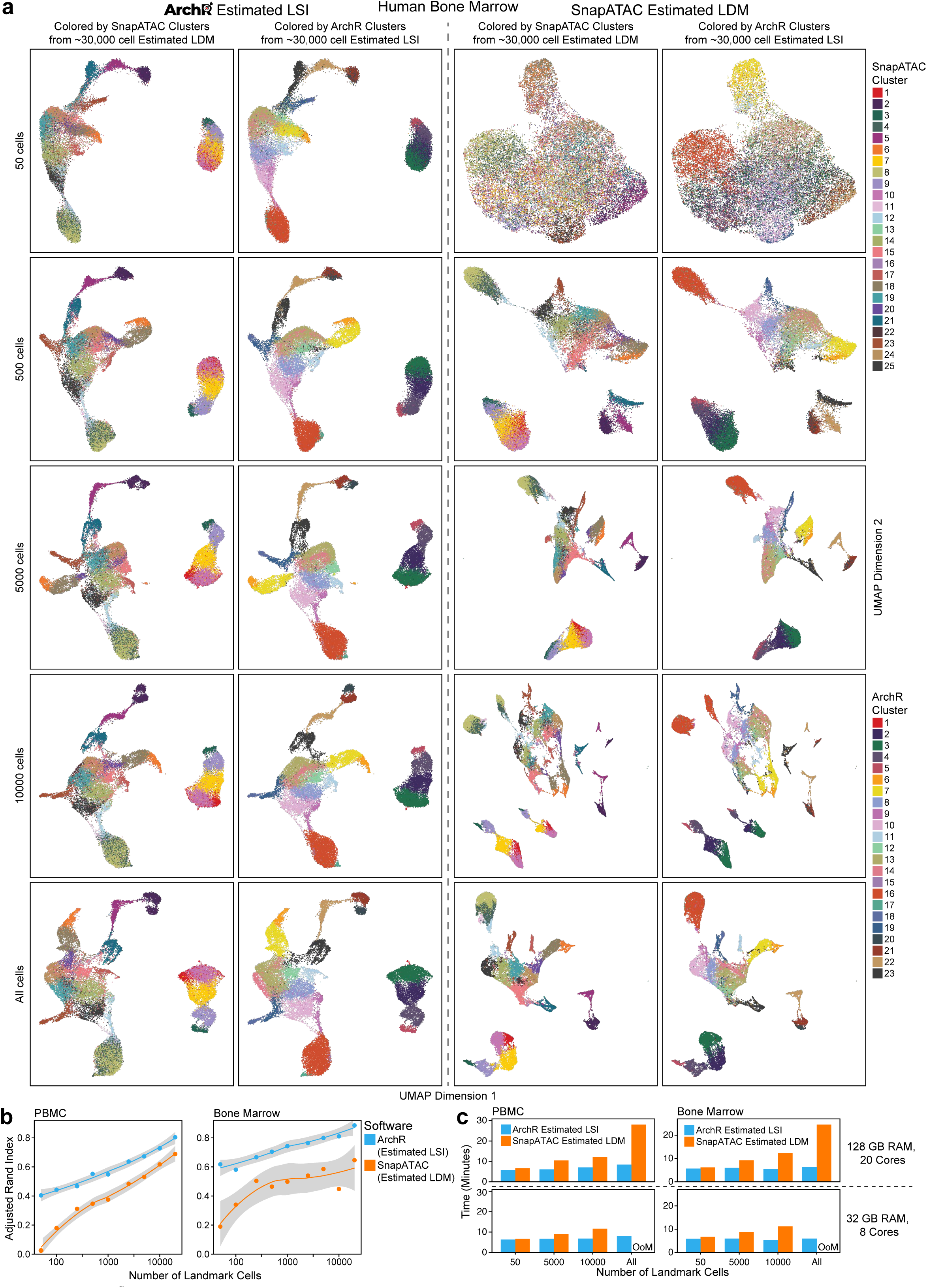
**a.** UMAPs of scATAC-seq data from ∼30,000 cells from the bone marrow cell dataset showing the results of dimensionality reduction from (left) estimated LSI with ArchR after doublet removal or (right) estimated LDM with SnapATAC. For each analytical case, a range of cell numbers is used for the landmark cell subset (top to bottom). Within each analytical case, two UMAPs are presented, colored by the clusters identified without estimation from (left) ArchR or (right) SnapATAC. **b.** Comparison of clustering fidelity based on adjusted Rand index in ArchR by estimated LSI or in SnapATAC by estimated LDM across multiple landmark subset sizes. **c.** Benchmarking of run time for ArchR estimated LSI and SnapATAC estimated LDM for ∼30,000 cells from (left) the PBMCs dataset and (right) the bone marrow cell dataset for (top) 128 GB of RAM with 20 cores and (bottom) 32 GB of RAM with 8 cores.

**Supplementary Fig. 9.**
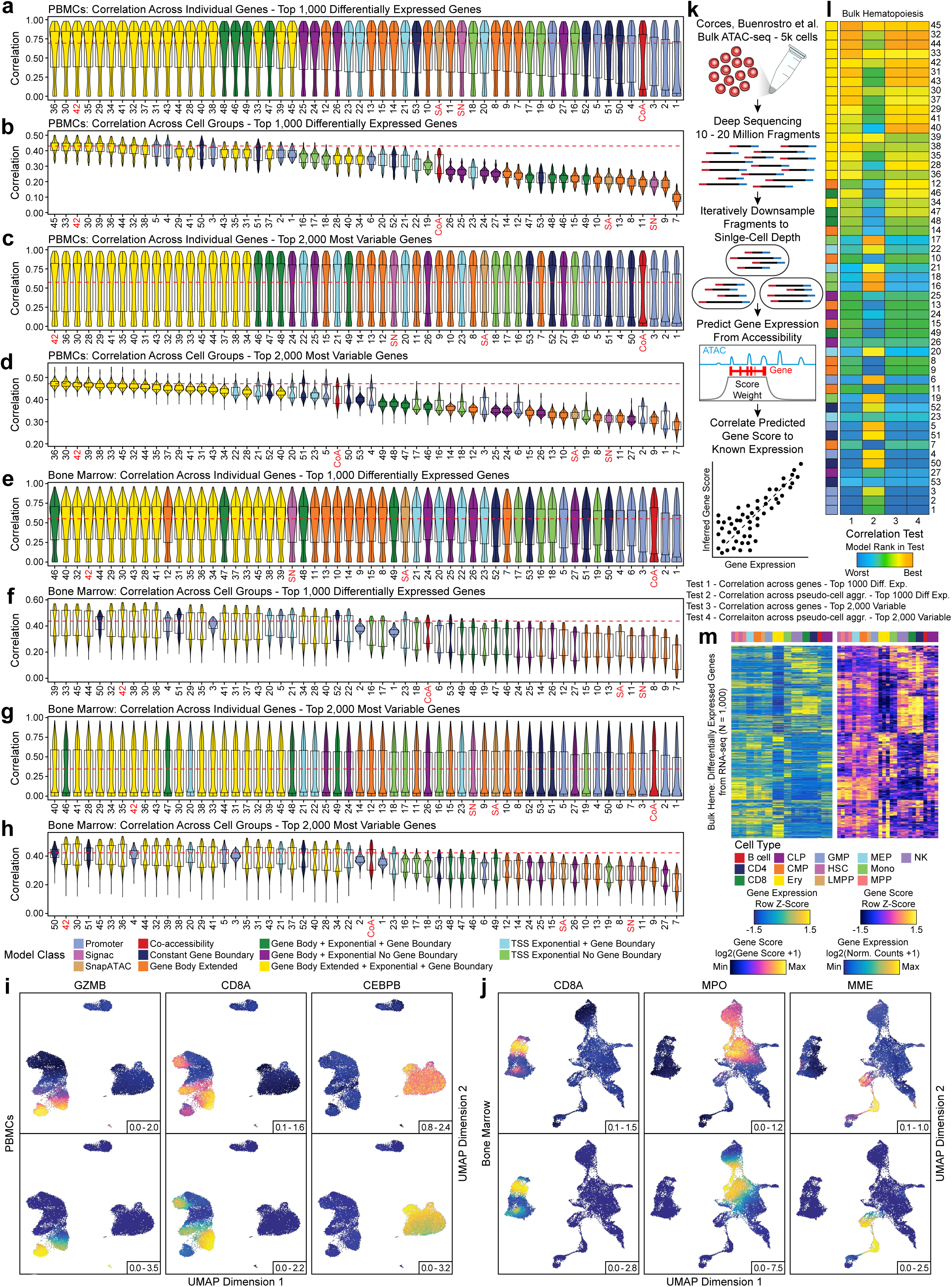
a-h. Distribution of Pearson correlations of inferred gene score and aligned gene expression for (**a,c,e,g**) each gene or (**b,d,f,h**) each cell group across groups of 100 cells (N = 500 groups). Distributions are either presented for (**a,b,e,f**) the top 1,000 differentially expressed genes or (**c,d,g,h**) the top 2,000 most variable genes for each of the 56 gene score models tested. In each plot, the red dotted line represents the median value of the best-performing model. Violin plots represent the smoothed density of the distribution of the data. In box plots, the lower whisker is the lowest value greater than the 25% quantile minus 1.5 times the interquartile range, the lower hinge is the 25% quantile, the middle is the median, the upper hinge is the 75% quantile and the upper whisker is the largest value less than the 75% quantile plus 1.5 times the interquartile range. SA, SnapATAC; SN, Signac; CoA, Co-accessibility. **i-j.** UMAPs of scATAC-seq data from (**i**) cells from the PBMCs dataset (N = 27,845 cells) or (j) cells from the bone marrow cell dataset (N = 26,748 cells) colored by (top) inferred gene scores or (bottom) gene expression for several marker genes. **k.** Schematic illustrating the methodology used to assess the accuracy of inferred gene scores based on orthogonal matched bulk ATAC-seq and bulk RNA-seq data of various sorted hematopoietic cell types. **l.** Heatmaps summarizing the accuracy (measured by Pearson correlation) across 56 gene score models for both the top 1,000 differentially expressed and top 2,000 variable genes for bulk ATAC-seq and RNA-seq data from sorted hematopoietic cell types. Each heatmap entry is colored by the model rank in the given correlation test as described below the heatmap. The model class is indicated to the left of each heatmap by color. SA, SnapATAC; SN, Signac; CoA, Co-accessibility. **m.** Heatmaps of (left) gene expression or (right) gene scores for the top 1,000 differentially expressed genes (selected from bulk RNA-seq) across all cell types from the matched bulk ATAC-seq and RNA-seq data.

**Supplementary Fig. 10.**
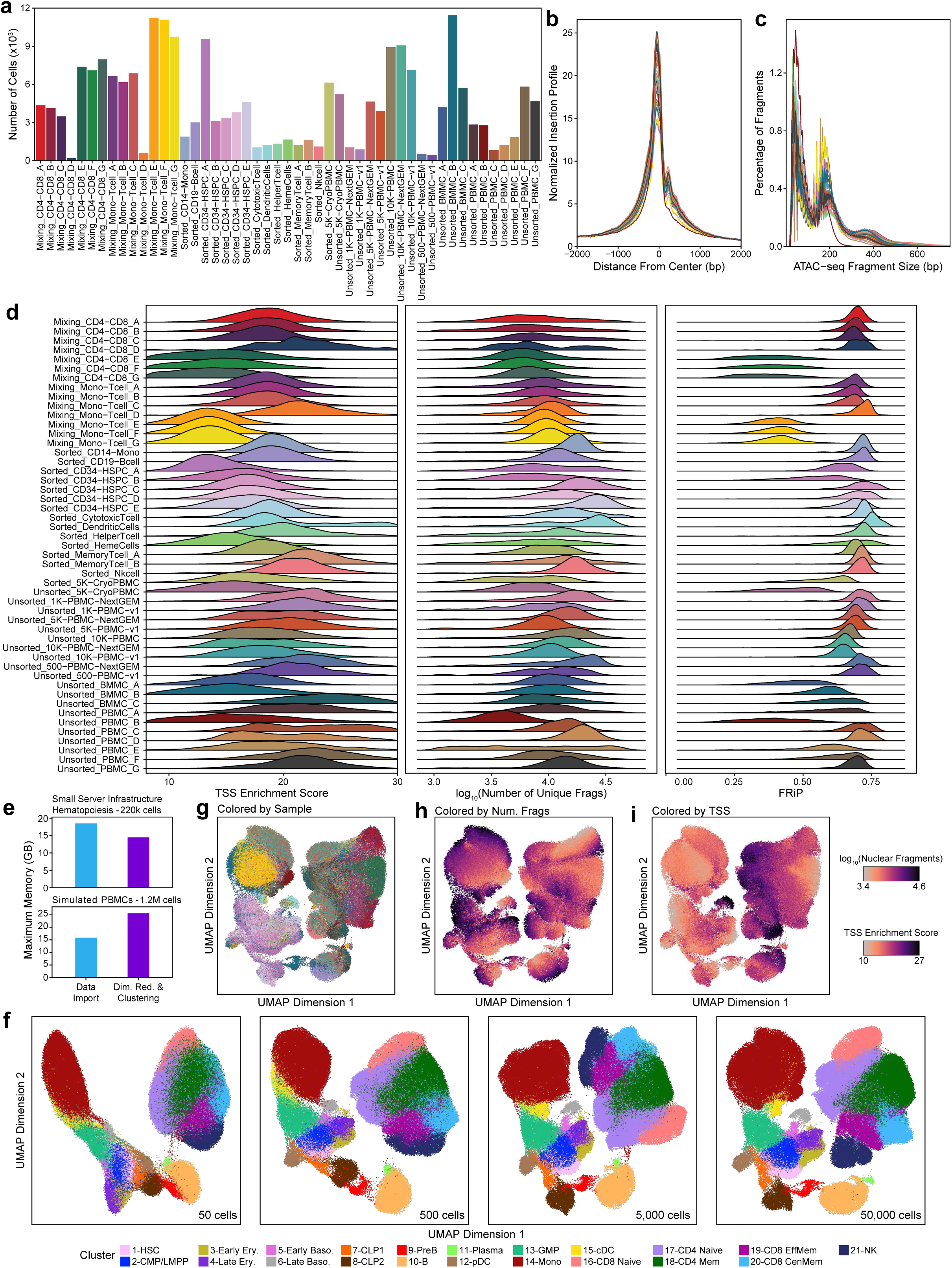
**a.** Bar plot showing the number of cells passing ArchR QC thresholds from each of the immune cell scATAC-seq datasets used for the ∼220k cell hematopoiesis dataset. **b-c.** Aggregate (**b**) TSS insertion profiles centered at all TSS regions or (**c**) fragment size distributions for the cells passing ArchR QC thresholds for each sample in the hematopoiesis dataset. Line color represents the sample from the dataset as indicated in **Supplementary Figure 10a**. **d.** Summary of quality control information for each cell from the hematopoiesis dataset. The distribution of (left) TSS enrichment scores, (middle) the number of unique nuclear fragments, and (right) the fraction of reads in peak regions (FRiP) are shown for each single cell passing filter. **e.** Benchmarking of peak memory usage for analysis of (top) the ∼220,000 cells from the hematopoiesis dataset and (bottom) ∼1,200,000 simulated PBMCs using a computational infrastructure with 32 GB of RAM and 8 cores with an HP Lustre file storage system. **f.** UMAPs of scATAC-seq data derived from estimated LSI of the hematopoiesis dataset using different numbers of landmark cells. These UMAPs are colored by the clusters identified from the 25,000-cell estimated LSI shown in Figure 3b. **g-i.** UMAPs of scATAC-seq data as shown in Figure 3b, colored by (**g**) the different experimental samples (as shown in **Supplementary Figure 10a**), (**h**) the number of unique nuclear fragments, or (**i**) the per-cell TSS enrichment score.

**Supplementary Fig. 11.**
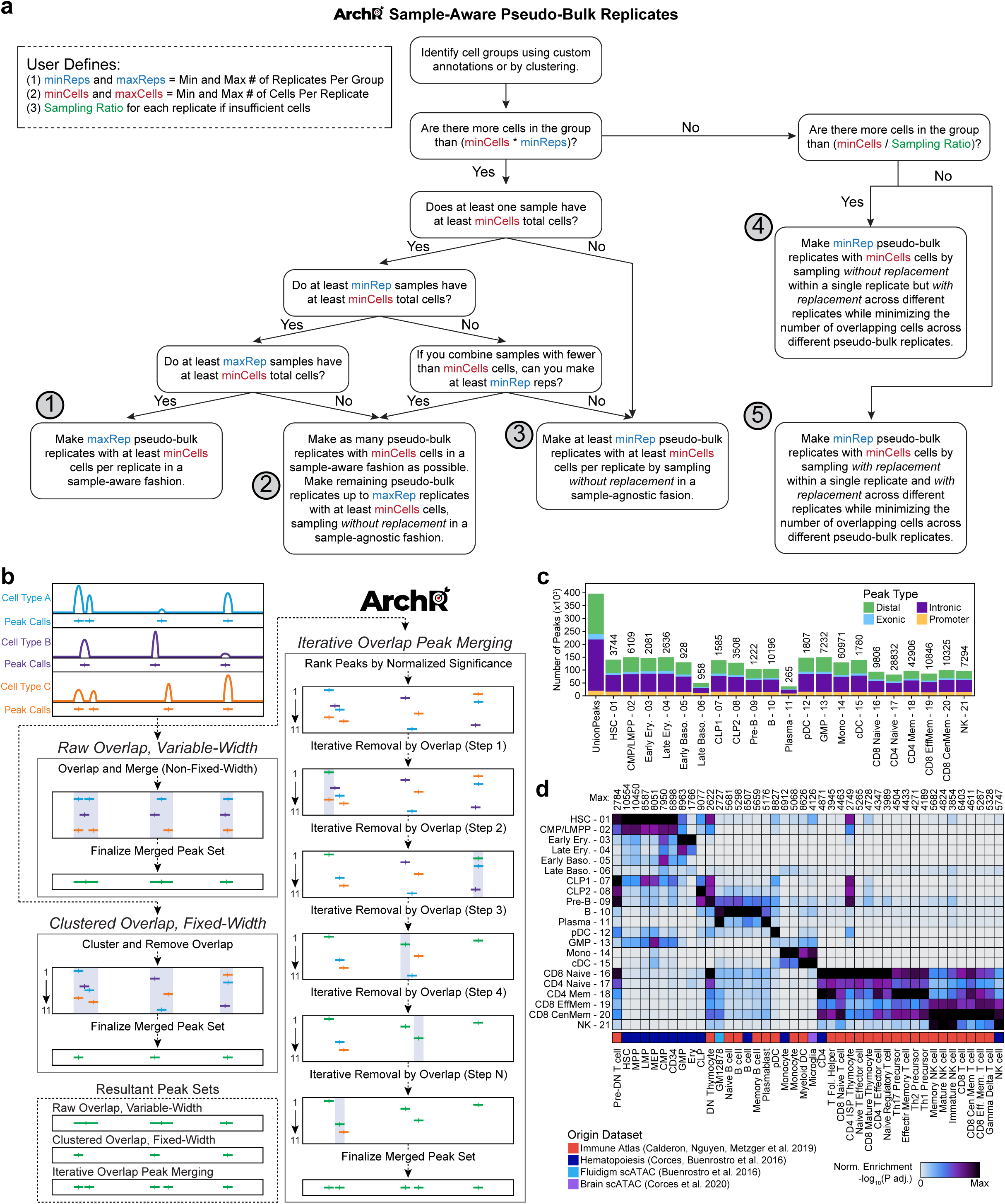
**a.** Schematic for the generation of sample-aware pseudo-bulk replicates in ArchR for downstream analyses. Briefly, for each cell grouping (in most cases identified by clusters), cells are split per sample of origin. Next, for each cell grouping these sample-aware cell groups are tested for being larger than a specified minimum number of cells to create a specified minimum number of sample-aware replicates. If these requirements are not met with a simple splitting, ArchR accounts for each different case by using sub-sampling approaches (see methods). **b.** Schematic for iterative overlap peak merging in ArchR to identify non-overlapping fixed-width peaks. Briefly, peaks (peak summits that are extended to yield fixed-width peaks) are called per sample and then ranked by significance. Next, for all peaks across multiple samples, the peak with the highest significance is kept. Peaks directly overlapping this most-significant peak are discarded and then this procedure is repeated until all peaks have either been kept or discarded, thus converging upon a non-overlapping fixed-width peak set. **c.** Bar plot showing the number of final peaks identified across all clusters (“Union Peaks”) and within each cluster from the hematopoiesis dataset. Bars are colored by peak annotation relative to a supplied gene set. **d.** Heatmap of hypergeometric enrichment testing the overlap of curated peak sets from previously published bulk ATAC-seq data (provided by ArchR) with the marker peak sets identified for each cluster in the hematopoiesis dataset in Figure 3c.

**Supplementary Fig. 12.**
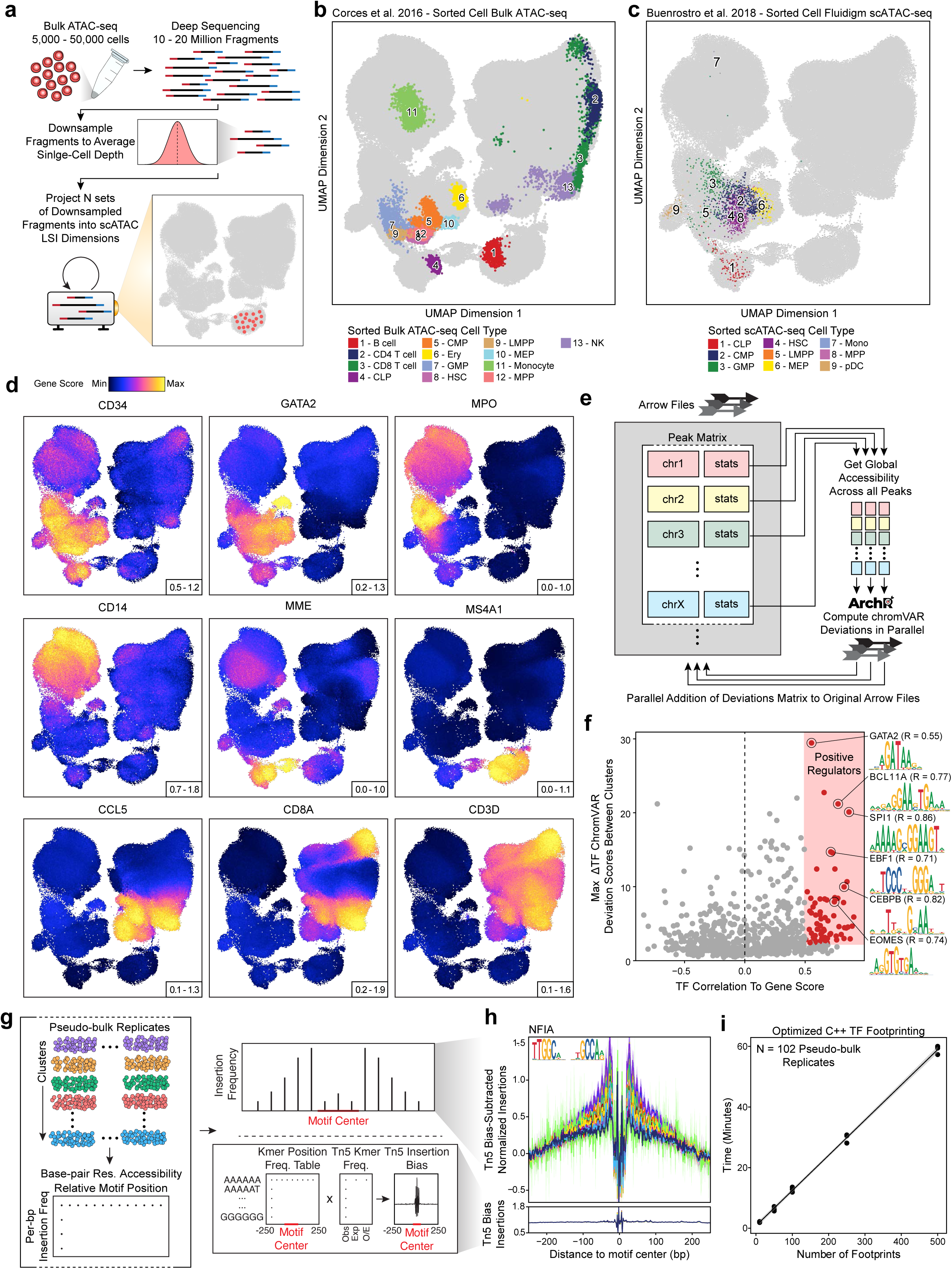
**a.** Schematic for the projection of bulk ATAC-seq data into an existing single-cell embedding using LSI projection. Briefly, bulk ATAC-seq data is deeply sequenced (10-20 million fragments), down sampled to a fragment number corresponding to the average single-cell experiment, and LSI-projected into the single-cell subspace. **b.** LSI projection of bulk ATAC-seq data from diverse hematopoietic cell types into the scATAC-seq embedding of the hematopoiesis dataset. **c-d.** UMAP of scATAC-seq data from the hematopoiesis dataset (N = 215,031 cells) colored by (**c**) sorted cells processed with the Fluidigm C1 system or (**d**) inferred gene scores for marker genes of hematopoietic cells. **e.** Schematic of the scalable chromVAR method implemented in ArchR. Briefly, ArchR computes global accessibility within each peak and then computes chromVAR deviations for each sample independently. This design facilitates large-scale chromVAR analysis with minimal memory usage for massive-scale scATAC-seq datasets. **f.** Dot plot showing the identification of positive TF regulators through correlation of chromVAR TF deviation scores and inferred gene scores in cell groups (Correlation > 0.5 and Deviation Difference in the top 50^th^ percentile). These TFs were additionally filtered by the maximum observed deviation score difference observed across each cluster average. This additional filter removes TFs that are correlated but do not have large accessibility changes in hematopoiesis. **g.** Schematic of TF footprinting with Tn5 bias correction in ArchR. Briefly, base-pair resolution insertion coverage files from sample-aware pseudo-bulk replicates are used to compute the insertion frequency around each motif for each replicate. For each motif, the total observed k-mers relative to the motif center per bp are identified. This k-mer position frequency table can then be multiplied by the individual sample Tn5 k-mer frequencies to compute the Tn5 insertion bias per replicate. **h.** TF footprint for the NFIA motif. Lines are colored by cluster identity from the hematopoiesis dataset shown in Figure 3b. **i.** Benchmarking of run time for TF footprinting with ArchR for the 102 sample-aware pseudo-bulk replicates from the hematopoiesis dataset.

**Supplementary Fig. 13.**
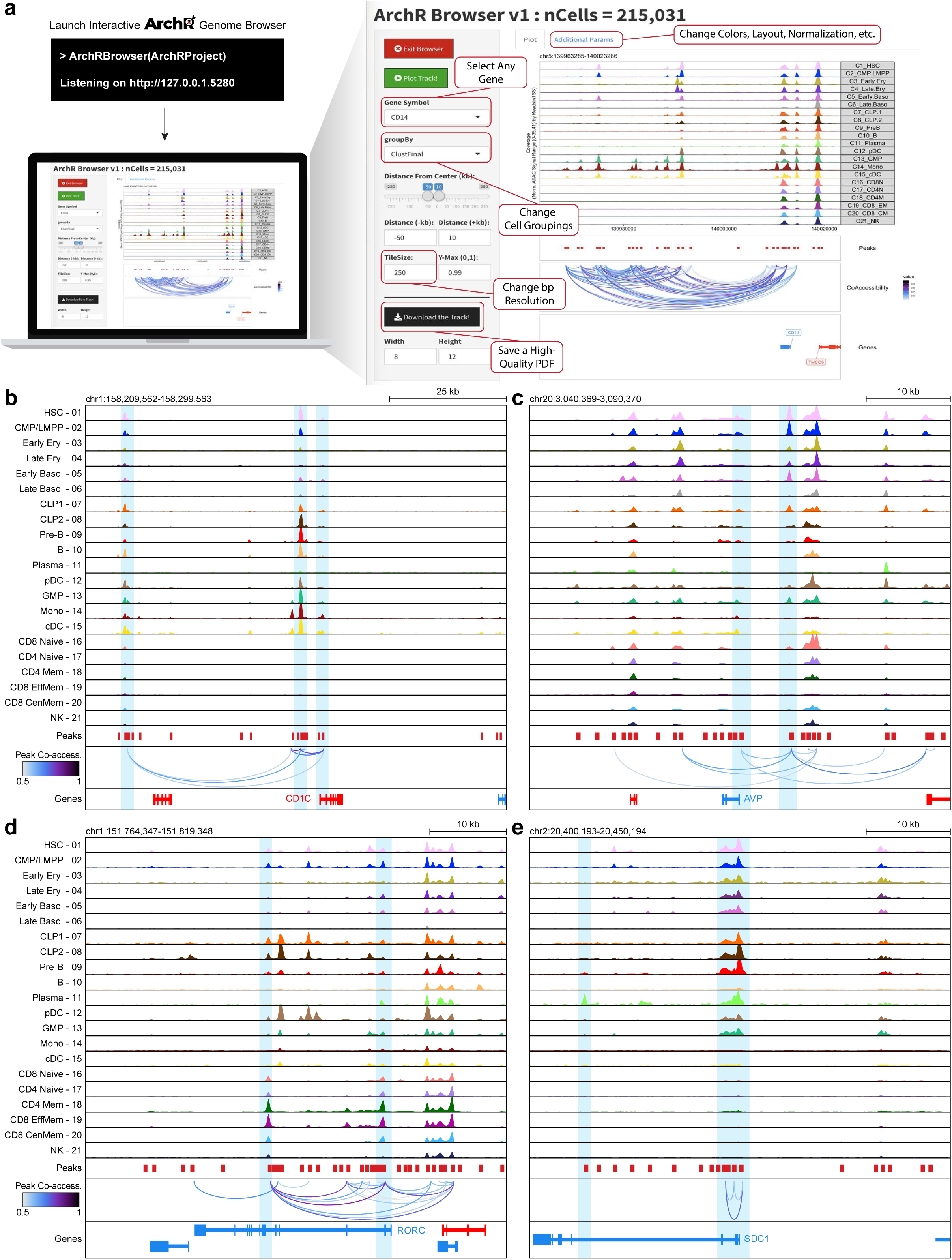
**a.** Schematic of the ArchR integrative genome browser. Briefly, the ArchR integrative browser is launched with a single command into an interactive Shiny session. From there, users can select any gene to visualize the accessibility genome track. Additionally, users can change cell groupings, resolution, layout and more with an intuitive user interface. Lastly, users can supply custom feature regions (such as peak sets) or looping/linkage sets (such as peak co-accessibility). **b-e.** Genome accessibility track visualization of marker genes with peak co-accessibility for (**b**) the *CD1C* locus (chr1:158,209,562-158,299,563), (**c**) the *AVP* locus (chr20:3,040,369-3,090,370), (**d**) the *RORC* locus (chr1:151,764,347-151,819,348), and (**e**) the *SDC1* locus (chr2:20,400,193-20,450,194).

**Supplementary Fig. 14.**
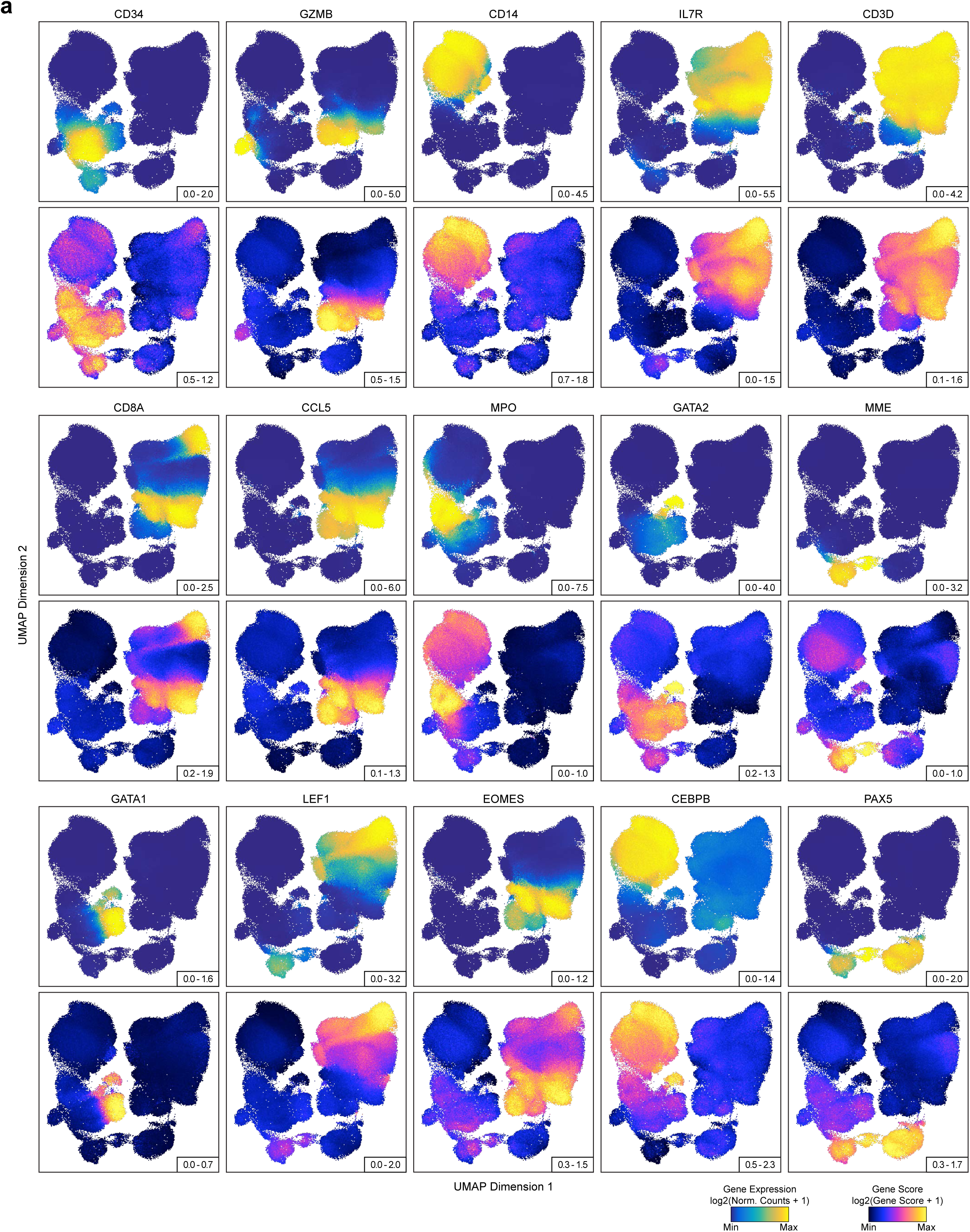
**a.** Side-by-side UMAPs for the hematopoiesis dataset cells colored by (top) gene expression (log2(Normalized Counts + 1)) from scRNA-seq alignment or (bottom) inferred gene scores (log2(Gene Score + 1)) from gene score Model 42 (see Figure 2c) for key immune marker genes.

**Supplementary Fig. 15.**
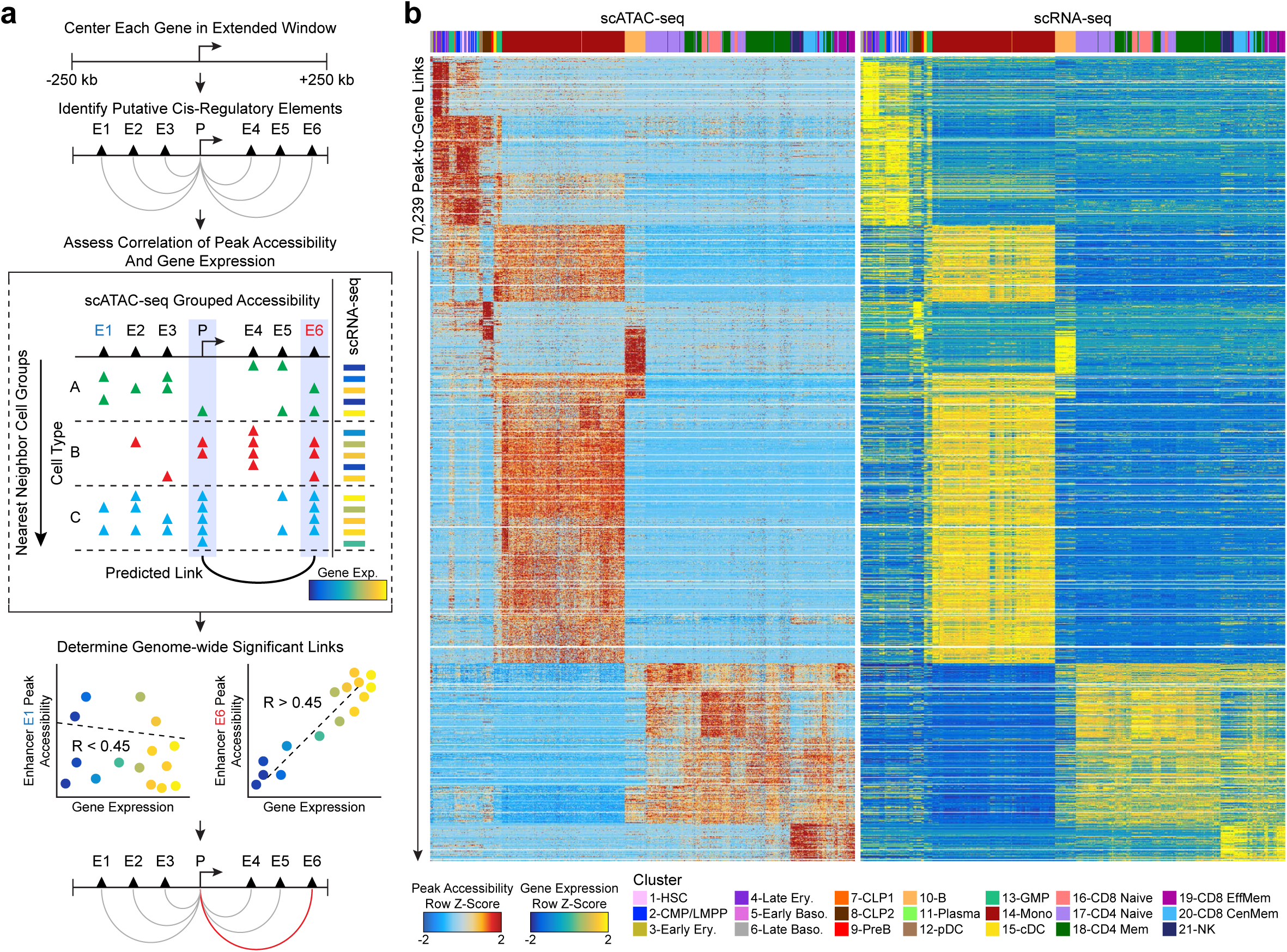
**a.** Schematic of identification of peak-to-gene links with ArchR. First, all combinations of peak-to-gene linkages are identified. Second, the peak accessibility and gene expression for cell groups are calculated. Finally, all potential peak-to-gene linkages are tested and significant links (R > 0.45 and FDR < 0.1) are kept. **b.** Heatmap of 70,239 peak-to-gene links identified across the hematopoiesis dataset with ArchR.

## Supplementary Tables

**Supplementary Table 1. scATAC-seq Data Sets**

This table contains information about each scATAC-seq data set used in this study including QC statistics, scATAC platform and source.

**Supplementary Table 2. scATAC-seq Benchmarking Results**

This table contains information corresponding to benchmarking results of Signac, SnapATAC and ArchR for the benchmarking data sets used in this study. Information such as run time and maximum memory usage are present in this table.

**Supplementary Table 3. Gene Score Models**

This table contains information for each of the Gene Score models used in Figure 2. Descriptions of each model are provided in this table.

**Supplementary Table 4. Positive Hematopoietic Regulators**

This table contains information for the identification of positively correlated Hematopoietic TFs. Information such as Pearson correlation, linkage statistics and motif are located in this table.

**Supplementary Table 5. Hematopoiesis Peak To Gene Linkages**

This table contains information corresponding to the peak to gene linkages in Hematopoiesis. Information such as peak coordinate, gene coordinate and Pearson correlation can be found in this table.

## References

1. Buenrostro, J. D. et al. Single-cell chromatin accessibility reveals principles of regulatory variation. Nature 523, 486–490 (2015).

2. Cusanovich, D. A. et al. Multiplex single cell profiling of chromatin accessibility by combinatorial cellular indexing. Science 348, 910–914 (2015).

3. Cusanovich, D. A. et al. The cis-regulatory dynamics of embryonic development at single-cell resolution. Nature 555, 538–542 (2018).

4. Buenrostro, J. D. et al. Integrated Single-Cell Analysis Maps the Continuous Regulatory Landscape of Human Hematopoietic Differentiation. Cell 173, 1535–1548.e16 (2018).

5. Cusanovich, D. A. et al. A Single-Cell Atlas of In Vivo Mammalian Chromatin Accessibility. Cell 174, 1309–1324.e18 (2018).

6. Satpathy, A. T. et al. Massively parallel single-cell chromatin landscapes of human immune cell development and intratumoral T cell exhaustion. Nat. Biotechnol. 37, 925–936 (2019).

7. Granja, J. M. et al. Single-cell multiomic analysis identifies regulatory programs in mixed-phenotype acute leukemia. Nat. Biotechnol. 37, 1458–1465 (2019).

8. Lareau, C. A. et al. Droplet-based combinatorial indexing for massive-scale single-cell chromatin accessibility. Nat. Biotechnol. 37, 916–924 (2019).

9. Chen, H. et al. Assessment of computational methods for the analysis of single-cell ATAC-seq data. Genome Biol. 20, 241 (2019).

10. Fang, R. et al. Fast and Accurate Clustering of Single Cell Epigenomes Reveals *Cis* -Regulatory Elements in Rare Cell Types. BioRxiv (2019). doi:10.1101/615179

11. Stuart, T. et al. Comprehensive Integration of Single-Cell Data. Cell 177, 1888–1902.e21 (2019).

12. McGinnis, C. S., Murrow, L. M. & Gartner, Z. J. DoubletFinder: Doublet Detection in Single-Cell RNA Sequencing Data Using Artificial Nearest Neighbors. Cell Syst. 8, 329–337.e4 (2019).

13. Wolock, S. L., Lopez, R. & Klein, A. M. Scrublet: Computational Identification of Cell Doublets in Single-Cell Transcriptomic Data. Cell Syst. 8, 281–291.e9 (2019).

14. Kang, H. M. et al. Multiplexed droplet single-cell RNA-sequencing using natural genetic variation. Nat. Biotechnol. 36, 89–94 (2018).

15. Thurman, R. E. et al. The accessible chromatin landscape of the human genome. Nature 489, 75–82 (2012).

16. Andersson, R. & Sandelin, A. Determinants of enhancer and promoter activities of regulatory elements. Nat. Rev. Genet. 21, 71–87 (2020).

17. Arnosti, D. N. Analysis and function of transcriptional regulatory elements: insights from Drosophila. Annu Rev Entomol 48, 579–602 (2003).

18. Pliner, H. A. et al. Cicero Predicts cis-Regulatory DNA Interactions from Single-Cell Chromatin Accessibility Data. Mol. Cell 71, 858–871.e8 (2018).

19. Corces, M. R. et al. Lineage-specific and single-cell chromatin accessibility charts human hematopoiesis and leukemia evolution. Nat. Genet. 48, 1193–1203 (2016).

20. McInnes, L., Healy, J. & Melville, J. UMAP: Uniform Manifold Approximation and Projection for Dimension Reduction. arXiv (2018).

21. Corces, M. R. et al. The chromatin accessibility landscape of primary human cancers. Science 362, (2018).

22. Calderon, D. et al. Landscape of stimulation-responsive chromatin across diverse human immune cells. Nat. Genet. 51, 1494–1505 (2019).

23. Corces, M. R. et al. Single-cell epigenomic identification of inherited risk loci in Alzheimer’s and Parkinson’s disease. BioRxiv (2020). doi:10.1101/2020.01.06.896159

24. Mumbach, M. R. et al. Enhancer connectome in primary human cells identifies target genes of disease-associated DNA elements. Nat. Genet. 49, 1602–1612 (2017).

25. Satpathy, A. T. et al. Transcript-indexed ATAC-seq for precision immune profiling. Nat. Med. 24, 580–590 (2018).

26. Corces, M. R. et al. An improved ATAC-seq protocol reduces background and enables interrogation of frozen tissues. Nat. Methods 14, 959–962 (2017).

27. Schep, A. N., Wu, B., Buenrostro, J. D. & Greenleaf, W. J. chromVAR: inferring transcription-factor-associated accessibility from single-cell epigenomic data. Nat. Methods 14, 975–978 (2017).

28. Regev, A. et al. The human cell atlas. Elife 6, (2017).

29. Huber, W. et al. Orchestrating high-throughput genomic analysis with Bioconductor. Nat. Methods 12, 115–121 (2015).

30. Adey, A. et al. Rapid, low-input, low-bias construction of shotgun fragment libraries by high-density in vitro transposition. Genome Biol. 11, R119 (2010).

31. Buenrostro, J. D., Giresi, P. G., Zaba, L. C., Chang, H. Y. & Greenleaf, W. J. Transposition of native chromatin for fast and sensitive epigenomic profiling of open chromatin, DNA-binding proteins and nucleosome position. Nat. Methods 10, 1213–1218 (2013).

32. Korsunsky, I. et al. Fast, sensitive and accurate integration of single-cell data with Harmony. Nat. Methods 16, 1289–1296 (2019).

33. Lun, A. T. L., McCarthy, D. J. & Marioni, J. C. A step-by-step workflow for low-level analysis of single-cell RNA-seq data with Bioconductor. [version 2; peer review: 3 approved, 2 approved with reservations]. F1000Res. 5, 2122 (2016).

34. Fornes, O., et al. JASPAR 2020: update of the open-access database of transcription factor binding profiles. Nucleic Acids Res. 48, D87–D92 (2020).

35. Sheffield, N. C. & Bock, C. LOLA: enrichment analysis for genomic region sets and regulatory elements in R and Bioconductor. Bioinformatics 32, 587–589 (2016).

36. He, H. H. et al. Refined DNase-seq protocol and data analysis reveals intrinsic bias in transcription factor footprint identification. Nat. Methods 11, 73–78 (2014).

37. Baek, S., Goldstein, I. & Hager, G. L. Bivariate genomic footprinting detects changes in transcription factor activity. Cell Rep. 19, 1710–1722 (2017).

38. van Dijk, D. et al. Recovering Gene Interactions from Single-Cell Data Using Data Diffusion. Cell 174, 716–729.e27 (2018).

